# BARTharm: MRI Harmonization Using Image Quality Metrics and Bayesian Non-parametric

**DOI:** 10.1101/2025.06.04.657792

**Authors:** Emma Prevot, Dieter A. Häring, Laura Gaetano, Russell T. Shinohara, Chris C. Holmes, Thomas E. Nichols, Habib Ganjgahi

## Abstract

Image derived phenotypes (IDPs) harmonization from Magnetic Resonance Imaging (MRI) data is essential for reducing scanner-induced, non-biological variability and enabling accurate multi-site analysis. Existing methods like ComBat, while widely used, rely on linear assumptions and explicit scanner IDs - limitations that reduce their effectiveness in real-world scenarios involving complex scanner effects, non-linear biological variation, or anonymized data. We introduce `BARTharm`, a novel harmonization framework that uses Image Quality Metrics (IQMs) instead of Scanner IDs and models scanner and biological effects separately using Bayesian Additive Regression Trees (BART), allowing for flexible, data-driven adjustment of IDPs. Through extensive simulation studies, we demonstrate that IQMs provide a more informative and flexible representation of scanner-related variation than categorical Scanner IDs, enabling more accurate removal of non-biological effects. Leveraging this and its ability to model complex relationships, `BARTharm`, consistently outperforms ComBat across a range of challenging scenarios, including model misspecification and confounded scanner-biological relationships. Applied to real-world datasets, `BARTharm` successfully removes scanner-induced bias while preserving meaningful biological signals, resulting in stronger, more reliable associations with clinical outcomes. Overall, we find that `BARTharm` is a robust, data-driven improvement over traditional harmonization approaches, particularly suited for modern, large-scale neuroimaging studies.

## 1 Introduction

With the rise of collaborative data-sharing initiatives, such as large population neuroimaging cohorts and clinical consortia, multi-site data collection has become essential for large-scale studies due to logistical constraints and the need to capture geographic variability in subject populations (Laird, 2021; Van Horn & Toga, 2009). However, integrating data from multiple scanning sites introduces unwanted non-biological variability, which must be addressed to ensure accurate and reproducible analyses. In neuroimaging, a key challenge is inter-scanner biases, often referred to as scanner effects, which arise when subject data are acquired using different MRI scanners, field strengths, and site-specific acquisition protocols. These differences can lead to systematic differences in Image Derived Phenotypes (IDP) distributions across scanning sites due to factors such as scanner hardware differences, acquisition parameters, time-of-day effects, and other environmental influences (Clark et al., 2006; Goto et al., 2012; Han et al., 2006; Jovicich et al., 2006; Takao et al., 2011; Trefler et al., 2016). To address this, harmonization is crucial for removing non-biological variability while preserving true biological differences in IDPs. Without proper harmonization, inter-scanner variability introduces both bias and noise. Bias can lead to spurious associations when scanner effects confound biological variables, while noise reduces sensitivity to true effects. Harmonization is essential to correct for bias and reduce variability, ensuring valid and powerful neuroimaging analyses.

Harmonization techniques in neuroimaging can be broadly classified into prospective and retrospective approaches (Hu et al., 2023). Prospective harmonization is integrated into the study design, ensuring that all data are acquired under standardized conditions, typically by using the same scanner model, field strength, and scanning protocol across multiple sites. However, prospective harmonization is not an option when leveraging and combining already existing datasets, as standardization would have had to be implemented before data collection. Retrospective harmonization is instead applied post hoc to correct for scanner-induced biases using already collected data. Existing retrospective harmonization methodologies can be broadly categorized into statistical, deep learning, and normative modeling approaches, each offering distinct strategies for addressing scanner-induced biases in MRI data. Deep learning approaches, including GANs, autoencoders, and other network architectures, can operate at both the image level and the IDP level and typically learn mappings that transform raw or feature-level data from one scanner domain to another, reducing scanner-specific artifacts (Moyer et al., 2020; Sohn et al., 2015; Tang et al., 2019; Zhu et al., 2017). Normative modeling, on the other hand, situates individual scans within a reference population, providing individualized predictions that implicitly correct for site effects without directly adjusting the data (Bayer et al., 2022). However, both deep learning and normative modeling carry notable limitations: deep learning methods often require extensive training data covering the full range of scanner variability and still struggle to generalize to new sites or protocols (domain shift), while normative modeling relies on a reference population that may not capture all scanner and population variations, limiting its applicability to diverse or novel datasets. Statistical methods have gained a lot of popularity for their flexibility and relative simplicity and form the backbone of harmonization techniques. These often apply location-scale adjustments or latent factor modeling to disentangle biological signals from batch effects.

Among the statistical methods, ComBat is the most widely adopted retrospective harmonization technique in neuroimaging (Fortin et al., 2017). It decomposes each IDP’s mean into two linear components — one representing the biological (covariate-based) effect and the other capturing the non-biological (scanner-specific) effect. The non-biological component is modeled as both an additive (mean shift) and a multiplicative (variance scaling) term across imaging sites, with these scanner-specific parameters estimated using empirical Bayes methods (Fortin et al., 2017). Harmonization is performed by centering and scaling the data based on these estimates, followed by reintroducing the biological component, all ensuring that scanner-induced variability is removed while preserving meaningful biological signal. Several conceptual extensions to ComBat have also been developed. ComBat-GAM enables the preservation of non-linear biological covariate effects through the use of generalized additive models (GAM) (Pomponio et al., 2020); longitudinal ComBat refines the modeling of serial measurements by incorporating random effects (Beer et al., 2020); CovBat, extends ComBat by harmonizing not just means and variances but also the covariance structure of residuals (Chen et al., 2022), and DeepResBat introduces deep residual batch harmonization to account for differences in covariate distributions (An et al., 2024).

Despite its practical success (Acquitter et al., 2022; Barth et al., 2023; Campello et al., 2022; Dai et al., 2022; Ingalhalikar et al., 2021; Pagani et al., 2023; Radua et al., 2020), ComBat has notable limitations. First, it assumes scanner effects as well as biological covariates of interest influence IDPs in a strictly linear manner, which does not account for the complex, non-linear effects often observed in neuroimaging data. For example, brain structures follow intricate developmental and age-related changes that cannot always be captured with linear models (Fjell et al., 2010). Additionally, ComBat requires a sufficiently large sample size per scanner to ensure stable estimates, making it less reliable in studies where only a few subjects are scanned at each site. A further shortcoming of many existing harmonization methods, including ComBat, is the reliance on scanner or site identifiers to estimate non-biological variability, which can be problematic in anonymized datasets where such information is often suppressed to protect data privacy rights. Moreover, recent research has demonstrated that Image Quality Metrics (IQMs), which quantify intrinsic image properties such as contrast-to-noise ratio and spatial resolution, provide a richer representation of scanner effects than simple site labels (Esteban et al., 2017, 2019). Studies have shown that IQMs are strongly associated with MRI scanners and acquisition protocols, making them highly informative for harmonization (Esteban et al., 2017). For example, variations in grey matter and white matter contrast have been directly linked to scanner-dependent acquisition protocols (Tardif et al., 2009). Importantly, IQMs can be computed directly from individual MRI scans without requiring knowledge of scanner identity, making them particularly advantageous in anonymized datasets. While existing methods leveraging IQMs are limited, Neuroharmony (Garcia-Dias et al., 2020) represents a notable attempt in this direction. Neuroharmony trains a machine learning model to predict ComBat-based harmonization corrections using IQMs extracted with MRIQC, allowing for site-independent harmonization. However, as it fundamentally builds upon ComBat, it inherits the limitations of ComBat’s linear assumptions and reliance on precomputed batch effects.

To address these limitations, we propose BARTharm, a novel statistical harmonization method that leverages Bayesian Additive Regression Trees (BART) (H. A. Chipman et al., 2010) to separately model biological and scanner effects without assuming linearity. Unlike ComBat, BARTharm uses Image Quality Metrics (IQMs) instead of scanner IDs, enabling harmonization in datasets where scanner IDs are unavailable or those with minimal scanner-level data. By employing a non-parametric Bayesian approach, BARTharm flexibly captures complex, non-linear scanner-induced biases and preserves biological variation across IDPs from either single or multiple sites. We demonstrate BARTharm’s effectiveness under a range of challenging scenarios, including non-linear scanner effects, model misspecification from incomplete biological covariates, and strong correlations between biological and scanner variables. In all of these, we show BARTHarm has superior performance in identifying and removing latent unwanted scanner-induced variations, also drastically reducing False Positive Rate (FPR) and increasing power. By harnessing the rich information encoded in IQMs and employing a Bayesian non-parametric modeling approach, BARTharm represents a flexible, robust alternative to traditional harmonization techniques, improving statistical efficiency of multi-site neuroimaging studies.

The rest of the paper is organized as follows. Section 2 describes our proposed method, BARTharm, and compares it to ComBat’s modelling approach. In Section 3 we describe the simulation studies implemented to evaluate BARTharm’s performance and compare it to ComBat’s, while in Section 4 we present both simulation results as well as real-world application of BARTharm. Finally, we conclude with a discussion in Section 5.

## 2 Methods

We compare two different harmonization procedures: (1) Removal of site effects using ComBat (Johnson et al., 2007); (2) Removal of site effects using BARTharm, which is the novel method we propose. We use the following notation: let the outcome *y*_*jv*_ be the IDP for participant *j* and region *v*, for a total of *n* participants and *V* regions to be harmonized. Furthermore, let *x*_*j*_ be the 1 *× c* vector of covariates for participant *j*, for a total of *c* covariates. We can further divide *x*_*j*_ into *w*_*j*_ and *k*_*j*_ being respectively the biological covariates of interest and the IQMs covariates. We also introduce an index for the imaging site *s*, and one for the scan *i* to explain ComBat.

### 2.1 ComBat harmonization

ComBat is a widely used harmonization method that assumes scanner effects introduce both additive and multiplicative biases. It employs empirical Bayes to improve the estimation of site parameters, particularly in small sample sizes. The ComBat model assumes that the observed IDP measurements can be expressed as a linear combination of biological covariates and site effects, where the error term is modulated by additional site-specific scaling factors. Mathematically, the model can be expressed as:

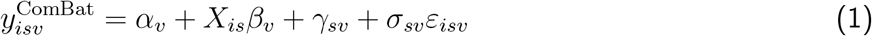

where *α*_*v*_ is the overall mean for IDP *v, X*_*is*_ is the design matrix for biological covariates, *β*_*v*_ is the corresponding IDP-specific regression coefficient vector, *γ*_*sv*_ and *σ*_*sv*_ represent the additive and multiplicative site effects, respectively, and *ε*_*isv*_ is a normally distributed error term.

ComBat harmonization adjusts for site effects by transforming the observed data into a harmonized version:

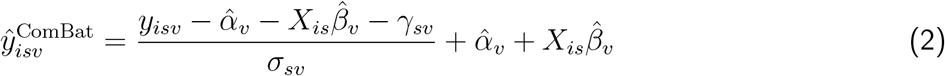

Despite its widespread adoption, we argue ComBat has notable limitations. The model assumes a linear site effect and requires scanner ID labels, which may not always be available in anonymized datasets. Furthermore, ComBat also struggles when there are only a handful of subjects from a scanner as it requires a statistically representative sample from each scanner included in the study. To address these challenges, we introduce BARTharm, a novel non-linear harmonization approach.

### 2.2 BARTharm harmonization

BARTharm adjusts the images for site effects using Bayesian Additive Regression Trees (BART) non-parametric modelling strategy (H. Chipman et al., 2006; H. A. Chipman et al., 2010).

#### 2.2.1 Bayesian Additive Regression Trees (BART)

We focus our discussion on continuous outcomes Bayesian Additive Regression Trees (BART) (H. Chipman et al., 2006; H. A. Chipman et al., 2010). We define our set of covariates *X* = *{x*_1_, ….*x*_*c*_*}* and the continuous outcome is *y*. A single regression tree represents the conditional expectation of *y* given *X* that is *E*[*y*|*X*]. We begin with a *root* node which is associated to a first splitting condition. If the condition is true for individual (*x*_*j*_) then follows path to the right, otherwise follows the path to the left. The path continues until the individual *j* reaches a *terminal* node, which is a node that is not split upon. Each terminal node has a parameter *µ* associated to it which is simply the value of *E*[*y*|*X*] for the individuals that went down that specific path. We denote by *T*_*k*_ the binary tree structure of the *k*^*th*^ tree, which is the set of binary decision rules; *M*_*k*_ is instead the collection of terminal nodes parameters of the tree, such that *M*_*k*_ = *{µ*_1,*k*_, …, *µ*_*b,k*_*}* where *b* is the total number of terminal nodes. One regression tree is formalised as *g*(*X*; *T*_*k*_, *M*_*k*_). Given an outcome *y*, BART tries to approximate a non parametric function *f* (*X*) as a sum of *m* regression trees, or a decision tree ensemble, such that

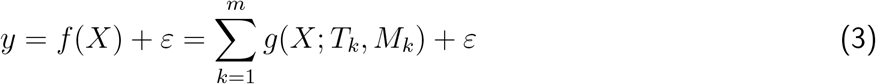

where 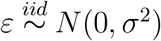. For simplicity we can also write *f ∼ BART*.

We complete the BART model specification by discussing the prior over all the parameters of the sum-of-trees model. The tree components (*T*_*k*_, *M*_*k*_) are independent of each other and of *σ*, and the terminal node parameters of every tree are independent. This simplifies the prior specifications to *p*(*T*_*k*_), *p*(*µ*_*kl*_|*T*_*k*_) and *p*(*σ*), where *µ*_*kl*_ *∈ M*_*k*_ is the set of terminal nodes for *k*^*th*^ tree. For *p*(*T*_*k*_), the trees are regularized to be shallow and weak: the probability of a split at node depth *d* is given by *α*(1 + *d*)^−*β*^, with typically *α* = 0.95 and *β* = 2 which strongly favors small trees, i.e., simpler models, such that the probability of a tree with 1, 2, 3, 4, and over 5 terminal nodes is 0.05, 0.55, 0.28, 0.09, and 0.03, respectively (Hill et al., 2020). For *p*(*µ*_*kl*_|*T*_*k*_) the terminal nodes parameters are drawn from 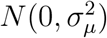 where *σ*_*µ*_ is set to ensure that the sum of trees *f* (*X*) lies within a plausible range based on the observed outcome. This prior effectively shrinks individual tree contributions and stabilizes the ensemble. Finally, the error variance *σ*^2^ is also given a conjugate inverse-chi-squared prior, calibrated using the data to ensure it spans a plausible range without being overly informative. The number of trees *m* is typically set to a default of 200, which provides strong predictive performance in most settings, with only minor degradation for larger values. This results in a model where each tree plays a modest role, and the ensemble captures the overall complexity of the function *f* (*X*).

The additive structure of BART enables it to flexibly capture both main effects and interactions by combining many shallow trees, each modeling different parts of the input space. Main effects arise when a tree splits on a single variable, while interactions emerge when a tree includes multiple variables in its split rules, allowing the effect of one variable to depend on another. Non-linearity is naturally captured through the piecewise-constant structure of each tree; their sum forms a smooth, adaptive approximation of complex functions. This ensemble approach allows BART to model varying levels of complexity across different regions of the covariate space. H. A. Chipman et al., 2010 showed through extensive empirical evaluation that BART consistently outperforms traditional models such as random forests, boosting, neural nets, and LASSO, often achieving the best predictive accuracy while providing well-calibrated uncertainty estimates.

#### 2.2.2 Adjusting for site effects with BARTharm

A recent extension of the standard BART model for treatment effect estimation is the Bayesian Causal Forest (BCF) framework (Hahn et al., 2018, 2020). The authors show that in the presence of strong confounding, standard nonlinear models, including BART, can yield biased causal estimates. This is especially problematic when the variation in the outcome due to prognostic covariates differs substantially from the variation due to treatment effects. In such cases, standard regularization may inadvertently shrink treatment effects toward zero. To address this, BCF decomposes the response surface into two components: one for the prognostic effect of the covariates and one for the treatment effect, each with its own BART prior.

In our work, we adapt this BCF framework for harmonization by leveraging a similar decomposition to separately model biological signals and scanner-induced variability. We write:

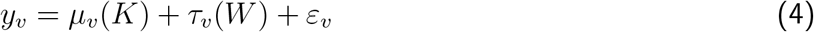

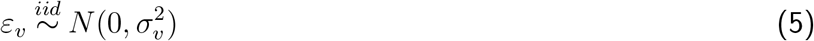

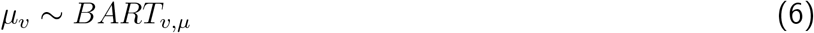

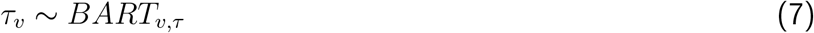

where *K* is the design matrix for the IQMs covariates, and *W* is the design matrix for the biological covariates. We separately model the effect of the biological covariates from the IQMs effect on the outcome *y*_*jv*_ for individual *j* and IDP *v*. Notice we have dropped here the site and scanner index as our model does not require any of this information. Moreover, by placing nonparametric BART priors on both the scanner-related component (*µ*) and the biological component (*τ*), BARTharm does more than just capture non-linear effects. It learns complex, high-order interactions and flexible functional relationships between covariates and the IDPs in a fully data-driven way. For the BART prior on the scanner effect, *BART*_*v,µ*_, we adopt the default specification of 200 trees with *α* = 0.95 and *β* = 2. For the one on biological effects *BART*_*v,τ*_ we use 100 trees, *β* = 3 and *η* = 0.25. This makes tree splitting considerably less likely, which is equivalent to shrinking more strongly toward homogeneous effects. Depending then on data properties, we might also decide to vary these hyperparameters to regularize more one effect.

It is important to note that although BARTharm is not explicitly formulated as a location-scale model, in its current implementation it assumes the differences in IDPs mean and variance across scanners can be explained by *µ*(*K*). By modeling scanner effects with IQMs using as a flexible, non-parametric BART prior, BARTharm can implicitly account for variance-related scanner effects, as long as the relevant variance structure is encoded in the IQMs.

After fitting our model, we obtain an estimate for the IQMs effect term 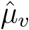, the biological effect 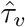, and the noise variance 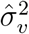. We evaluate the clean, harmonized data 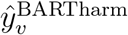 by removing the estimated scanner effect from the observed outcome:

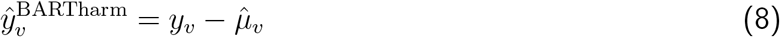

Unlike ComBat, which requires explicit scanner or site IDs, our model estimates scanner effects using IQMs which can be extracted from any MRI image regardless of scanner IDs, making it well-suited for anonymized datasets where identifiers are often unavailable or where only a single scan per site or scanner is available. This makes the method applicable to a broader range of real-world, multi-center studies without the limitations of site-based harmonization.

#### 2.2.3 Model fitting

In BART, model fitting and inference are performed using a custom Bayesian backfitting MCMC algorithm, which efficiently explores the high-dimensional posterior space of tree structures and terminal node values. At each iteration, the algorithm updates each tree *T*_*k*_ in turn, conditioning on the partial residuals, i.e., the portion of the outcome not yet explained by the other trees in the ensemble. This results in a residual vector *R*_*j*_ = *y* − *Σ*_*k≠j*_ *g*(*X*; *T*_*k*_, *M*_*k*_), which is treated as the response for updating the *j*-th tree. Each update consists of two steps: (1) proposing a new tree structure using a Metropolis-Hastings step (which may involve growing, pruning, or changing split rules), and (2) sampling new terminal node values from a conjugate normal prior, conditioned on the updated tree and residuals. This process yields a Markov chain that, after burn-in, generates samples from the posterior distribution over the space of sum-of-trees models.

To fit our `BARTharm` model, we adopt the approach proposed in the General BART framework (Tan & Roy, 2019), implementing a two-stage Gibbs sampler. This sampler alternately updates the scanner-related component *µ*_*v*_ and the biological component *τ*_*v*_, which are each modeled using independent BART ensembles, fit on the corresponding residuals.

Specifically, our Gibbs sampler alternates between updating the scanner-related effect *µ*_*v*_, the biological effect *τ*_*v*_, and the noise variance 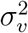, conditioned on the observed outcome *y*_*jv*_ and the relevant covariates. At each iteration, we draw from the following full conditional distributions:

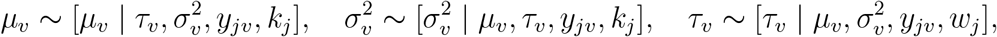

where *k*_*j*_ denotes the Image Quality Metrics (IQMs) and *w*_*j*_ denotes the biological covariates for subject *j*. This sampling scheme reflects the iterative structure of the Gibbs sampler, where the scanner-related and biological components are updated sequentially while conditioning on the current state of the other component and the observed data.

For the scanner effect *µ*_*v*_ update, we compute residuals *R*_*µ*_ = *y*_*jv*_ − *τ*_*v*_(*w*_*j*_), which represent the part of the outcome not explained by the biological component, and use them as the response to fit the scanner effect via BART. Conversely, when updating *τ*_*v*_, we compute residuals *R*_*τ*_ = *y*_*jv*_ − *µ*_*v*_(*k*_*j*_) and fit the biological contribution. This alternating procedure enables the model to disentangle scanner-induced variation from the true biological signal in a fully Bayesian, non-linear fashion.

To implement this in R we use the SoftBart R package (Linero, 2022; Ran & Bai, 2021) and create two BART *forest* object, one for *µ* and one for *τ*. In addition to enabling soft decision trees, SoftBart offers the notable advantage that researchers can integrate it into other R-based models without modifying the underlying C++ code, making it more accessible and adaptable for various applications (Linero, 2022; Ran & Bai, 2021). The SoftBART implementation requires the data to be normalised. Therefore, all biological and non-biological covariates are quantile normalized. We generally follow the literature to set the number of trees *m*, and the parameters for the splitting probability *α* and *β* (H. A. Chipman et al., 2010; Hahn et al., 2020).

##### Algorithm 1: BARTharm algorithm

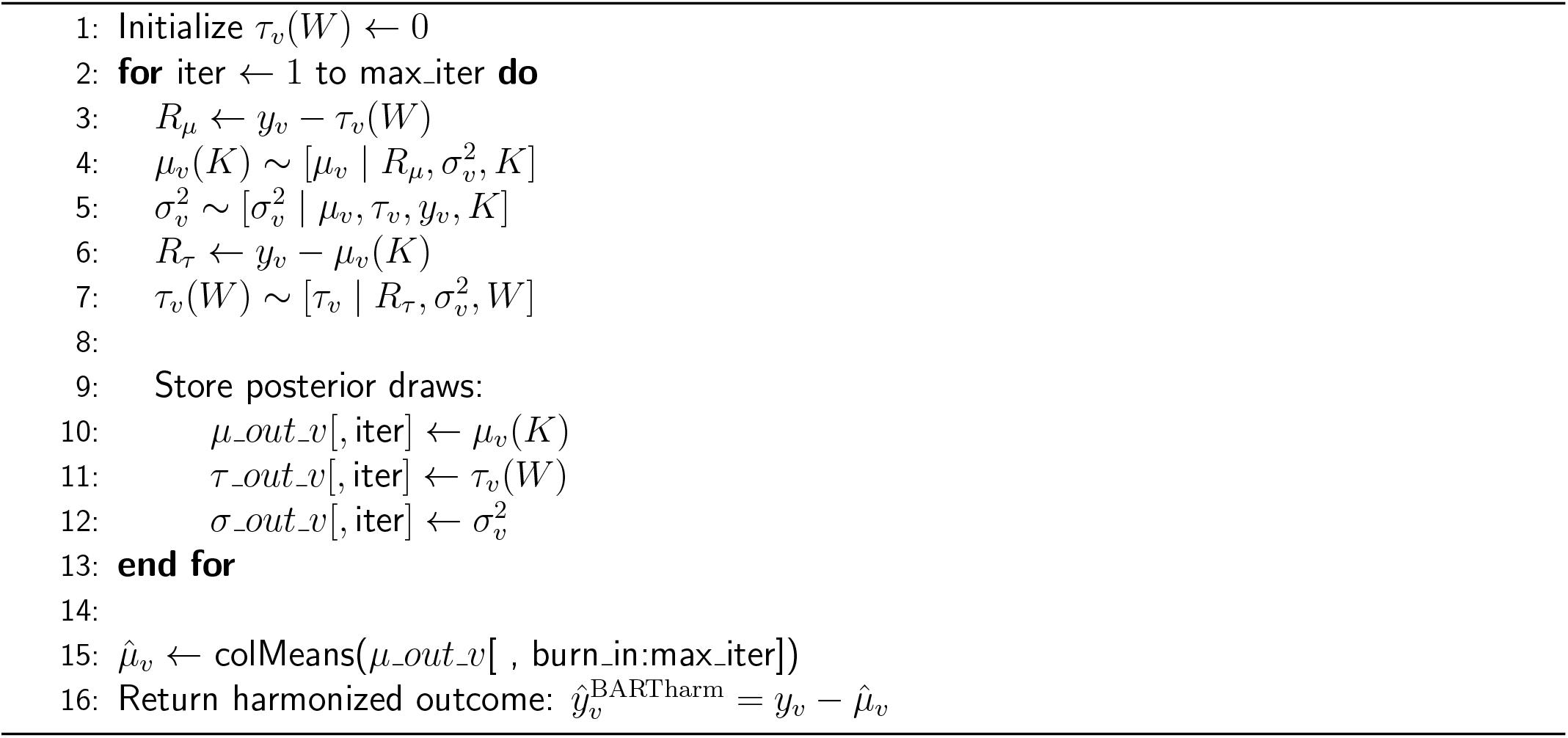

Algorithm 1 shows some of the key steps of the BARTharm algorithm. Once the for loop is completed, we obtain a full Markov chain for both *µ* and *τ* which we can use for a range of inferential tasks. We first assess the chain convergence via visual inspection of the posterior samples of the residual variance 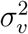 to define appropriate burn-in, thinning interval and number of iterations. We then retain the posterior draws after burn-in and average across those to get a point estimate for the biological effect 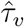 and the IQMs effect term 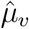, which we then use to evaluate the harmonized outcome 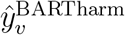. With the resulting sets of posterior draws we can also perform uncertainty quantification (via credible intervals), and functional summaries such as partial dependence plots, which reveal the marginal effect of covariates on either the estimated scanner effect term or the biological one. Additionally, BART supports model-free variable selection by tracking how frequently each covariate is used in tree splits across iterations.

This yields a data-driven estimate of variable importance that complements interpretability of the fitted harmonization surface.

Unlike empirical Bayes approaches, this fully Bayesian framework provides calibrated uncertainty quantification for each harmonized outcome. Similar advantages of full posterior inference were also demonstrated in recent work comparing empirical and fully Bayesian ComBat harmonization, where the latter was shown to better preserve biological signal and offer meaningful uncertainty estimates useful for downstream inference and augmentation tasks (Reynolds et al., 2023).

### 2.3 Data and preprocessing

#### 2.3.1 Extracting Image Quality Metrics

Image Quality Metrics (IQMs) are extracted using MRIQC, which is a tool to extract no-reference IQMs (image quality metrics) from structural (T1w and T2w), functional and diffusion MRI (magnetic resonance imaging) data (Esteban et al., 2017). The IQMs can be grouped in four broad categories, providing a vector of 56 features per anatomical image. Some measures characterize the impact of noise and/or evaluate the fitness of a noise model. A second family of measures uses information theory and prescribed masks to evaluate the spatial distribution of information. A third family of measures look for the presence and impact of particular artifacts. Specifically, the INU artifact, and the signal leakage due to rapid motion (e.g. eyes motion or blood vessel pulsation) are identified. Finally, some measures that do not fit within the previous categories characterise for instance the statistical properties of tissue distributions, volume overlap of tissues with respect to the volumes projected from MNI space, the sharpness/blurriness of the images, etc.

Importantly, BARTharm is not limited to a specific set of IQMs or to a particular extraction tool such as MRIQC. It can be flexibly applied to any collection of IQMs derived from any available quality assessment framework, provided they capture meaningful scanner-induced variability. Furthermore, BARTharm does not require a fixed number of IQMs as input; it can accommodate any number or subset of metrics, allowing users to tailor the harmonization process to the characteristics of their dataset and the quality indicators most relevant to their specific imaging protocol.

#### 2.3.2 NO.MS Dataset

The Novartis Oxford MS (NO.MS) dataset (Dahlke et al., 2021; Mallon et al., 2021) is the largest and most comprehensive clinical trial dataset in Multiple Sclerosis with more than 35,000 patients with up to 15 years of follow-up data. All trials were conducted in accordance with the provisions of the International Conference on Harmonization guidelines for Good Clinical Practice and the principles of the Declaration of Helsinki. All trial protocols were approved by an institutional review board or ethics committee and all patients or their legal representatives gave written informed consent before any trial-related procedures were performed. Data have been de-identified in a risk-based approach as reported by Mallon et al., 2021. In brief, identifiers (including facial features on scans) were either removed, generalised or modified to minimise the risk of re-identification.

For our real-data analysis we retained baseline observations for 5,386 individuals from the NO.MS. The outcomes we harmonize are 29 brain image-derived phenotypes (IDPs), including cortical gray matter volumes from deep subcortical regions and the cerebellum. Examples are Normalised Brain Volume (NBV), Thalamus, Parietal Lobe, Hippocampus, and we provide the complete list in the Supplementary Material Section 1.4.

We used the following covariates to model the biological contribution *τ* : sex, age, MS subtype, treatment type, Expanded Disability Status Scale (EDSS), Volume of T2 lesions, Number of gadolinium-enhancing lesions, number of relapses within one year before entering the trial, number of relapses within two years prior to entering the trial, and duration from the first symptoms. The scanner effect *µ* was instead modeled on all the IQMs extracted applying MRIQC on T1w images, except weighted summary statistics for each tissue distribution in the data.

## 3 Simulation studies

In this section, we describe a series of simulation studies designed to evaluate the performance of BARTharm and benchmark it against ComBat under varying conditions of scanner and biological variability. These experiments aim to assess each method’s ability to recover harmonized IDPs in the presence of linear or non-linear scanner and biological effects, and to test robustness to challenges like unobserved confounders and confounded site assignment. We distinguish between *fully synthetic* and *semi-synthetic* data. In the fully synthetic setting, both biological (e.g., age, sex) and non-biological (e.g., SNR, CNR) covariates and their effects are entirely simulated. In the semi-synthetic setting, we use real biological variables and IQMs from the NO.MS dataset, simulating only the outcomes based on user-defined effect structures. This hybrid setup allows us to evaluate harmonization accuracy under realistic yet controlled conditions.

On fully synthetic data we run four simulation studies (Simulation framework 1). These explore how well each method disentangles biological signal from scanner-induced noise under different combinations of linear and non-linear biological and scanner effects. The goal is to assess recovery of the true harmonized signal under a range of data-generating processes. On semi-synthetic data instead we run two simulation studies which reflect more realistic scenarios (Simulation framework 2). In the *misspecification* setting, the outcome is generated using all biological covariates, but each method is intentionally fit using only 50% of them. This mimics real-world conditions where some relevant biological covariates may be unobserved, poorly measured, or unavailable — leading to model misspecification and potential bias in harmonization. In the *correlation* setting, we introduce varying degrees of correlation between biological and scanner effects, simulating cases where scanner assignment is confounded with subject characteristics (e.g., one site scanning mostly patients, another mostly controls). This reflects the common challenge in retrospective studies of disentangling scanner-related variability from true biological differences.

### 3.0.1 Simulation setting 1: non-linear relationships and complex scanner effects

The goal of this simulation framework is to evaluate the performance of BARTharm and ComBat in recovering harmonized imaging-derived phenotypes (IDPs) under varying levels of complexity in both biological and scanner-related contributions. Specifically, we aim to test whether incorporating rich Image Quality Metrics (IQMs), rather than relying solely on Scanner IDs, improves harmonization, particularly in settings that go beyond the linear assumptions of ComBat. We simulate fully synthetic data to allow complete control over the data-generating process, enabling us to isolate and manipulate the functional forms of both the biological and scanner effects. When the biological signal is simulated linearly, the setup serves as a baseline where ComBat is expected to perform well. In this case, we compare its performance to BARTharm under both linear and non-linear scanner effects, to assess whether IQMs offer a more informative and effective basis for harmonization than Scanner ID alone. When the biological signal is instead simulated non-linearly, representing a more realistic scenario, we again introduce either linear or non-linear scanner effects. This setting allows us to demonstrate how BARTharm enables more accurate recovery of the harmonized signal by capturing non-linearities in both biological and scanner components.

To simulate these scenarios, we first define the properties of either 2 or 10 different MRI scanners by varying resolution, noise level, signal-to-noise ratio (SNR), and contrast-to-noise ratio (CNR). Scanner resolution is categorized into low, medium, and high, with noise levels inversely related to the scanner ID. SNR and CNR values are derived from uniform distributions, adjusted by noise level, to ensure realistic scanner-induced variability. To simulate subject-level data for 1,000 individuals, variables such as age, sex, and scanner ID are generated. Scanner IDs are randomly assigned to ensure no correlation with biological covariates. Two continuous biological features, independently drawn from uniform distributions, represent true biological signals unaffected by scanner effects. The outcome is composed of two components: a biological term *Y*_bio_ and a scanner-related term *Y*_scanner_, following the decomposition *Y*_sim_ = *Y*_bio_ + *Y*_scanner_ + *ε*, where *ε* is Gaussian noise with a proportional variance of 5-10% of (*Y*_bio_ + *Y*_scanner_). The baseline, biological, outcome *Y*_bio_ for each subject is modeled as a function of sex, age, and biological features, incorporating both main and interaction effects with different regression coefficients *β*_bio_. Scanner effects *Y*_scanner_ are then added to the biological outcome with regression coefficients *β*_scanner_, shared by all scanners, ensuring that resolution, noise level, SNR, and CNR influence the final measurement in a structured manner. Higher scanner resolution decreases the outcome, while higher noise levels increase it, with interaction effects between resolution and noise further modulating these relationships. Both biological and scanner contributions can be generated either linearly or non-linearly, depending on the experimental condition. Mathematical details including the specific regression coefficients used in the linear settings and the functional forms used to simulate non-linear effects are provided in the Supplementary Material at Section 1.2.

To evaluate the risk of false discoveries introduced by harmonization, we also randomly assign a binary group indicator to each individual, which we include as biological covariate. This variable is completely unassociated with the outcome, allowing us to test whether a harmonization method mistakenly amplifies spurious group differences (false positives). In contrast, to assess the sensitivity to detect true effects, we evaluate statistical power using a biological covariate with a known, non-zero effect on the outcome—in our case, sex. This covariate consistently induces a meaningful group difference, allowing us to measure how reliably harmonization methods preserve true signal and facilitate detection of real associations in downstream analyses.

Figure 1 provides an example of simulated data where scanner-related differences are statistically significant. The boxplots compare the median outcome at each site before and after introducing scanner noise; although the differences are not extensive, they provide a realistic multi-site scenario. We can also observe some differences across the spread of the distributions. As previously discussed, although BARTharm is not explicitly formulated as a location-scale model, we assume that the BART prior on the scanner effect component enables it to flexibly capture both location and scale variation, if such variation is explained by the IQMs. Therefore, we did not limit the simulated scanner effects to just location shifts, but they can also influence the scale (variance) of the measurements. We can test this formally for each realization of the data using Bartlett’s test for homogeneity of variances, which assesses whether the variance of the outcome differs significantly across scanners. A significant p-value from this test indicates that scanner-induced heteroskedasticity is present, which justifies the need for harmonization methods that account not only for mean shifts but also for differences in variance across sites.

**Figure 1:**
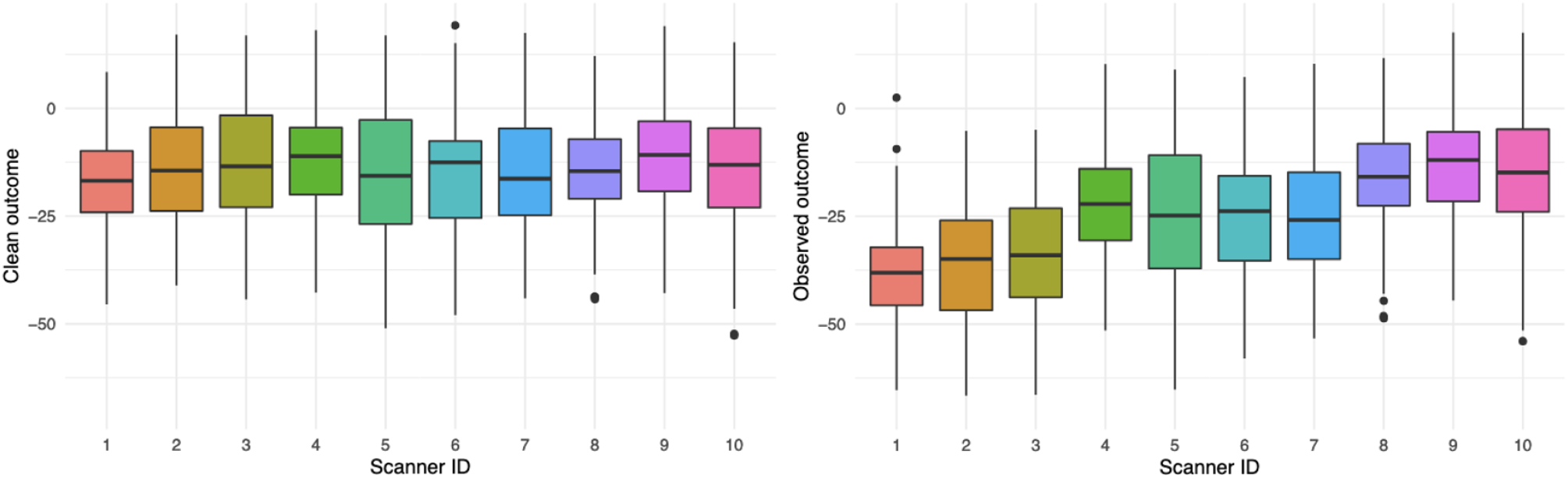
Distribution of simulated outcomes across ten scanners from one realization of Simulation setting 1. The left panel shows the clean outcome distribution (*Y*_*clean*_ = *Y*_*bio*_ + *ϵ*, while the right panel illustrates the same data after scanner-specific noise (*Y*_*scanner*_) has been introduced, which is the final simulated outcome *Y*_*sim*_. Each box plot summarizes the median (black horizontal line), interquartile range (colored box), and the 5–95% range (whiskers). Outliers beyond the 5–95% interval are shown as black dots. We observe that scanner-induced noise introduces substantial shifts and variability across scanners, highlighting the presence of systematic scanner effects

### 3.0.2 Simulation setting 2: Model misspecification and correlation

The second simulation framework is designed to assess the robustness of harmonization methods under more realistic conditions, using a semi-synthetic setup. Specifically, the goal is to evaluate whether methods like BARTharm and ComBat can accurately recover the true biological signal when faced with (i) incomplete biological information (model misspecification), and (ii) confounding between scanner assignment and biological variables (correlation). In this setup, we use real biological covariates and IQMs from the NO.MS dataset, extracting a subset of 1500 observations, and simulate scanner IDs and outcomes. To simulate Scanner ID, we apply k-means clustering to the IQM covariates and assign each subject to one of 5 clusters, which serves as proxy for the ID. The outcome is composed of two components: a biological term *Y*_bio_ and a scanner-related term *Y*_scanner_, following the decomposition *Y*_sim_ = *Y*_bio_ + *Y*_scanner_ + *ε*, where *ε* is Gaussian noise. To simulate *Y*_bio_, we sample a vector of regression coefficients *β*_bio_ *∼ 𝒩* (0, 4), which are then kept fixed across all scanners to ensure that the biological signal is independent of site. The biological outcome *Y*_bio_ is computed as a linear combination of these coefficients and the real biological covariates. Thus, *Y*_bio_ = *X*_bio_*β*_bio_, representing a linear biological signal. For the scanner component *Y*_scanner_, we introduce scanner-specific variation through a set of regression coefficients 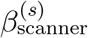 for each scanner *s* = 1, …, 5, sampled independently from a normal distribution. These coefficients are applied to the real IQM covariates such that 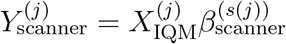, where *s*(*j*) denotes the scanner ID for subject *j*. This ensures individualized scanner contributions that reflect site-dependent biases in IQM distributions. As with the biological component, the scanner effects are simulated linearly in this setup. The final simulated outcome *Y*_sim_ combines both components with additive Gaussian noise, where the noise standard deviation is scaled relative to the signal magnitude. This semi-synthetic design provides a realistic yet controlled environment to assess each method’s ability to recover the harmonized signal and remove scanner effects under misspecification and correlation conditions.

For the *correlation* experiments, this framework is then extended to introduce a controlled correlation between biological (*Y*_bio_) and scanner (*Y*_scanner_) components, specifically 0.30, 0.50, and 0.75 correlation, while maintaining scanner-specific effects. This is done through the simulated regression coefficients in a way that allows us to control the correlation level. For the *misspecification* experiments, we instead used the semi-synthetic dataset without any correlation between the biological and scanner contributions. However, when fitting BARTharm and ComBat we only used 50% of the biological covariates, choosing them randomly at every realization.

We again introduce a randomly assigned group indicator, included as a biological covariate, to evaluate whether harmonization methods inadvertently induce spurious group differences. However, in this framework, we also simulate a non-zero correlation between the group and the IQM covariate that has, on average, the strongest contribution to the scanner effect component *Y*_scanner_ in each realisation. Hence, we expect to find a scanner-induced group bias in the simulated outcome *Y*_sim_, but not in the the true underlying biological outcome *Y*_clean_. This setup mimics a more challenging and realistic scenario where group assignment is partially confounded with scanner-related variability and highlights why harmonization is essential: to eliminate scanner-driven spurious associations that would otherwise be falsely detected in downstream analyses. As before, we assess sensitivity to detecting true biological effects using the sex covariate, which induces a known, non-zero impact on the outcome. Full mathematical details are provided in the Supplementary Material at Section 1.3.

#### 3.1 Performance evaluation

In all experiments, across both Simulation setting 1 and 2, we ran BARTharm Gibbs chain for 5,000 iterations, discarding the first 2,000 for burn-in, and we confirmed convergence by looking at the residual variance *σ*^2^ trace plots for a random subset of realizations. We configure the biological response forest *τ* and the scanner effect forest *µ* as described in Section 2.2.3. We simulate 1,000 realizations of the datasets and average the results across these simulations. Each realization undergoes a quality control check to ensure data integrity. To evaluate site-level differences, we apply ANOVA to determine whether the median simulated outcome varies significantly across the ten sites. If a significant difference is detected, we retain the realization; otherwise, it is discarded. Additionally, for the correlation experiments in the second simulation framework, we check that the simulated correlation is the desired one *±*0.05 as it can slightly vary across the different realizations.

We evaluate BARTharm and ComBat harmonization and compare their performance using different evaluation metrics. The first is the Root Mean Squared Error (RMSE) between the harmonized outcome *Y*_harm_ and the true, clean, outcome *Y*_clean_ = *Y*_bio_ + *ϵ*. RMSE captures the average deviation from the true outcome, with lower values indicating better harmonization accuracy. The second metric is BIO Corr, which refers to the Pearson correlation between *Y*_harm_ and *Y*_clean_. A BIO Corr close to 1 means that the harmonized data preserves the underlying biological variability, which is essential for meaningful downstream analysis. The third metric, IQMs Corr, measures the correlation between *Y*_harm_ and scanner-induced contribution *Y*_scanner_. A low IQMs Corr value, ideally near zero, indicates successful removal of scanner-related bias.

We then tested whether the two harmonization methods inadvertently introduce spurious group effects or fail to detect true ones. To do this, we computed two key metrics: the False Positive Rate (FPR) and statistical Power. FPR measures the proportion of times a significant group difference is detected when no true effect exists, allowing us to assess whether harmonization inflates false positives. Power, in contrast, measures the ability to correctly detect a real group difference when one is present. We assessed FPR using the simulated randomly assigned binary group indicator, which has no association with the outcome, and Power using the sex covariate, which induces a known effect. Both variables were included as covariates when fitting BARTharm and ComBat, allowing us to evaluate each method’s ability to preserve true biological signals while avoiding the introduction of spurious ones.

When interpreting the observed False Positive Rate (FPR) in our simulation experiments, it is important to account for the variability that arises from the finite number of simulation replicates. Under the null hypothesis, the number of spurious group effects detected across *N* simulations follows a binomial distribution with probability *p* = 0.05. Even if a method is well-calibrated, we do not expect the observed FPR to be exactly 0.05 in every experiment due to random variation. To determine whether an observed FPR is meaningfully higher than expected, we compute an upper confidence bound for 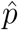, the sample estimate of the FPR, using a normal approximation to the binomial:

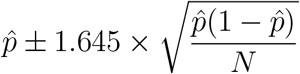

This expression gives the 95th percentile (one-sided) of the sampling distribution of the FPR under the null. Substituting 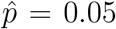 and *N* = 1000, we get an upper bound of approximately 0.06 and lower bound of 0.04. In other words, even a well-calibrated method can yield an observed FPR as high as 6% or as low as 4% just by chance. However, if the FPR exceeds the upper bound, it suggests the method is truly inflating Type I error, not just experiencing random fluctuation. Instead, an FPR below 4% indicates that the method is overly conservative, potentially sacrificing power to avoid false positives. Thus, in our simulation studies, FPRs above 0.06 are interpreted as significant inflation, while values below 0.04 signal excessive conservatism.

## 4 Results

### 4.0.1 Simulation results

Table 1 shows the results of Simulation setting 1, with fully-synthetic data and 10 simulated scanners, averaged across 1000 realizations. We observe that BARTharm (BARTh) consistently outperforms ComBat (ComB) across all configurations of biological (*µ*) and scanner-related (*τ*) effects. In the first row where both effects are linear ComBat performs reasonably well, as expected under its model assumptions. However, even in this favorable setting, ComBat fails to fully remove scanner-related variability, as indicated by an IQMs Corr of 0.174, which is substantially higher than the 0.007 observed for BARTharm. This highlights a key limitation: ComBat’s reliance on categorical Scanner IDs makes it unable to account for the finer-grained scanner variability captured by continuous IQMs. The same trend is evident in the second row, where the biological effects remain linear, but scanner effects are non-linear. Here, ComBat’s performance drops sharply, with BIO Corr falling to 0.381, IQMs Corr rises to 0.816, and RMSE increasing to 1.07. In contrast, BARTharm, which models scanner effects non-linearly using IQMs, maintains high biological correlation (0.999) and achieves substantially lower RMSE (0.65). In the bottom two rows, where biological effects are non-linear, ComBat’s limitations become even more apparent. Regardless of whether scanner effects are linear or non-linear, ComBat fails to capture the complex structure of the biological signal, with BIO Corr values around 0.74 and RMSE above 0.5. On the other hand, BARTharm retains near-perfect correlation with the true biological signal (0.999) and achieves substantially lower RMSE (0.24 and 0.23, respectively). Additionally, IQMs Corr values remain near-zero for BARTharm, confirming that it continues to suppress scanner-induced variability, even under the most challenging non-linear scenarios. Across all four settings, we highlight the robustness of BARTharm and the limitations of ComBat in capturing and removing complex scanner effects, particularly when the underlying relationships deviate from linearity.

**Table 1:**
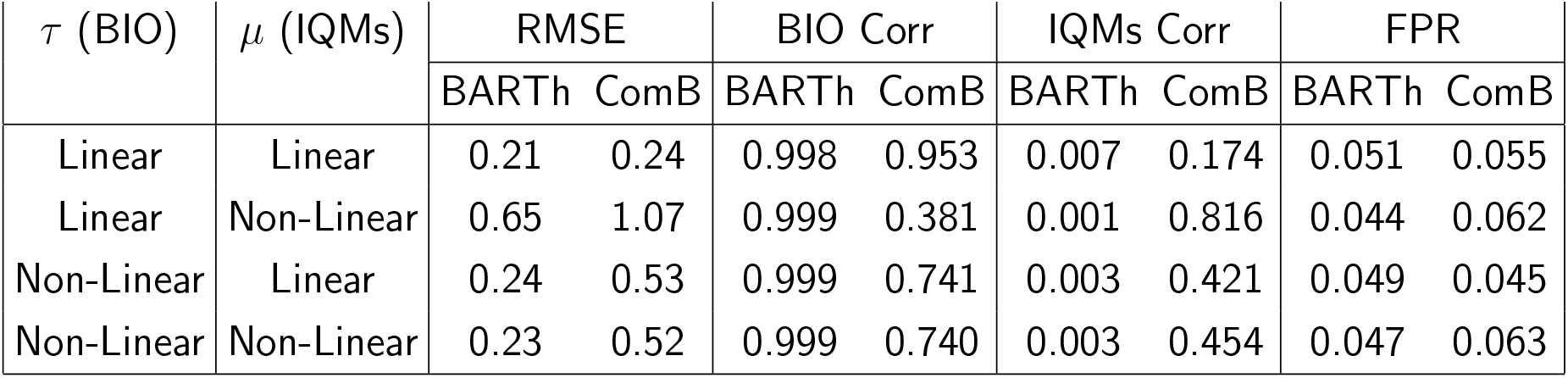
Comparison of BARTharm (BARTh) and ComBat (ComB) under different combinations of linear and non-linear biological (*τ*) and scanner (*µ*) effects, with 10 simulated scanners. RMSE indicates accuracy in recovering the true, harmonized, outcome, BIO Corr measures alignment with the true biological signal, IQMs Corr captures residual scanner effects, FPR denotes the false positive rate in detecting a group effect. Lower RMSE, higher BIO Corr, and lower IQMs Corr collectively signify more effective harmonization; an FPR above 0.06 is a statistically significant evidence of inflation, values below 0.04 signal excessive conservatism. BARTharm consistently outperforms ComBat across all settings, showing better accuracy, stronger biological signal recovery, and greater reduction of scanner-induced artifacts, particularly in scenarios with non-linear effects.

From Table 1 we can also observe that when both biological and scanner effects are linear, the FPR is well-calibrated for both methods, with values close to the nominal 0.05 level (0.051 for BARTharm and 0.055 for ComBat). However, in scenarios involving increased complexity, particularly when the IQM (scanner) effect is non-linear, ComBat exhibits slightly inflated FPRs, reaching 0.062 and 0.064 in the two most complex configurations. These values exceed the one-sided 95% confidence upper bound of 0.06, providing statistical evidence that ComBat may introduce spurious group effects in such settings. In contrast, BARTharm maintains FPRs within the expected range across all scenarios, confirming its ability to avoid false discoveries even under challenging, non-linear conditions. Notably, both methods maintain 100% power across all settings, indicating strong sensitivity to detect true effects when present.

Table 3 in the Supplementary Material instead shows the results obtained when simulating only two 2 scanners. The results are broadly consistent with those from the main simulation (Table 1), confirming that BARTharm consistently outperforms ComBat. Especially under non-linear data-generating processes, ComBat fails to adequately remove scanner-induced variation, as evidenced by high IQMs correlation and lower BIO correlation. In contrast, BARTharm maintains low residual scanner correlation and near-perfect recovery of the biological signal across all settings.

Finally, from Figure 1, we observed notable scale differences across scanners. To formally assess this, we applied Bartlett’s test for homogeneity of variances on each realization of every simulation setting (linear/non-linear *τ/µ*. We found differences in variances in all realizations and in at least 50% of the realizations per setting, the scanner-specific variances differed significantly, indicating the presence of significant scale effects. Despite this, BARTharm consistently outperformed ComBat, even though ComBat is an explicit location-scale harmonization model. This may suggests that BARTharm is capable of implicitly accounting for scanner-induced scale heterogeneity, due to the flexible modeling structure of scanner effects.

Table 2 shows the results from Simulation setting 2, averaged across 1000 realizations. Unlike the fully synthetic setting, here we use semi-synthetic data described in Section 3.0.2. The first row represents the *model misspecification* scenario, where the simulated outcome is generated using all biological covariates, but both BARTharm and ComBat are fit using only a randomly selected 50% of those covariates ^1^. This mimics a common real-world situation where key biological variables may be unobserved, poorly measured, or omitted from the model. In this setting, both methods experience some degradation in performance, but BARTharm remains more accurate and robust. It achieves a lower RMSE (2.17 vs. 2.25), higher biological correlation (0.95 vs. 0.78), and substantially better suppression of scanner-related variability (IQMs Corr = 0.03 vs. 0.23) compared to ComBat. These results indicate that BARTharm is better equipped to handle incomplete covariate information, maintaining higher fidelity to the true biological signal while minimizing residual scanner effects.

**Table 2:**
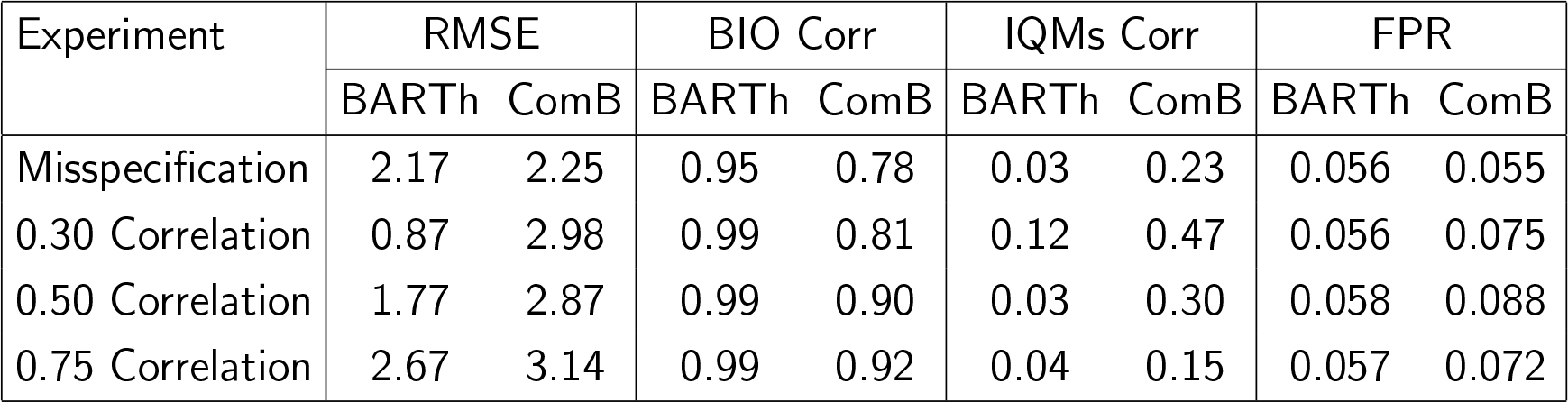
Performance comparison of BARTharm (BARTh) and ComBat (ComB) under model misspecification and varying correlation between biological and scanner effects. RMSE indicates accuracy in recovering the true, harmonized, outcome, BIO Corr reflects alignment with the true biological signal, IQMs Corr measures residual scanner influence, and FPR denotes the false positive rate in detecting a group effect. Lower RMSE, higher BIO Corr, and lower IQMs Corr collectively signify more effective harmonization; an FPR above 0.06 is a statistically significant evidence of inflation, values below 0.04 signal excessive conservatism. BARTh consistently outperforms ComBat, demonstrating greater robustness to model misspecification and correlated scanner-biological effects. It achieves lower RMSE, stronger recovery of the biological signal, and more effective removal of scanner-induced confounding across all tested conditions. Additionally, ComBat exhibits strong signs of FPR inflation.

The remaining rows evaluate performance under increasing levels of correlation between biological and scanner effects (0.30, 0.50, and 0.75), which is a realistic scenario in multi-site studies where certain scanners are disproportionately used for specific populations (e.g., patients vs. controls). As this correlation increases, the confounding becomes more severe. Given the complexity of these scenarios, perfect performance is not expected; however, these results highlight BARTharm’s superior resilience, making it a more reliable harmonization method under challenging conditions. More specifically, BARTharm’s performance is always superior to ComBat’s, maintaining near-perfect BIO Corr, very low IQM Corr, and smaller RMSE. While both methods are impacted by increasing correlation, as expected in any confounded scenario, BARTharm’s ability to use IQMs and non-linear modeling allows it to more effectively separate scanner-related noise from true biological signal. Additionally, BARTharm successfully mitigates scanner-driven spurious associations, maintaining FPR consistently within the expected confidence bounds (around 0.04–0.06) across all scenarios. In contrast, ComBat exhibits substantial FPR inflation when biological and scanner effects are correlated. For instance, ComBat’s FPR rises to 0.075 at 0.30 correlation and further to 0.088 at 0.50 correlation. This indicates that ComBat fails to fully remove scanner-induced bias under confounded settings, leading to an increased risk of false discoveries. Importantly, both methods maintain high Power, showing that their ability to detect true effects remains strong; however, only BARTharm preserves statistical validity by controlling false positives in challenging, realistic scenarios.

#### 4.1 Real Data analysis on NO.MS

We run our real data analysis on the NO.MS dataset. To protect patient privacy, the dataset was anonymized and does not include scanner or site identifiers, which is a common problem when integrating sensitive healthcare data from multiple sources, such as different clinical trials or imaging repositories, into a central data lake. This underscores the advantage of BARTharm’s ability to perform harmonization without relying on scanner IDs. For the same reason, we are unable to apply ComBat to NO.MS. We apply BARTharm to harmonize the 29 brain image-derived phenotypes (IDPs) described in Section 2.3.2, using the biological and IQMs covariates also described in that section. For the two BART forest, we use 200 trees for the biological response *τ* and 50 trees for the scanner response *µ*. Moreover, we set hyperparameters to more heavily penalise tree depth in the scanner response. These settings are different from what was described in Section 2.2.3. That is because we found the biological and IQMs covariates to be very heavily correlated and both highly predictive of the outcome. Therefore, we want to allow more flexibility in the biological response and more heavily regularise the scanner surface. We run the Gibbs chain for 100,000 iterations, with a thinning of 10 and burn-in of 2,500.

To assess whether variations in the imaging-derived phenotypes (IDPs) remain associated with scanner effects after harmonization, we visualize them as a function of the estimated scanner effect *µ*. Figure 2 presents a selection of representative IDPs. Before harmonization, there is a clear relationship between IDPs and scanner effects, driven by IQMs, as evidenced by strong trends in the plots. However, after applying BARTharm, this association is effectively removed, resulting in flattened lines and demonstrating that the method successfully eliminates scanner-induced biases. This can be tested formally by fitting a linear model with an interaction term between the scanner effect *µ* and a harmonization indicator. Specifically, we combine the observed and harmonized IDP in a single regression model and include an interaction term to estimate the change in slope before and after harmonization. A statistically significant interaction indicates a meaningful reduction in the association between the IDP and scanner effect due to harmonization. For all the 29 brain IDPs we get p-values ≪ 10^−16^, with post-harmonization slopes ranging from 0.05 to 0.2. This provides strong statistical evidence that BARTharm successfully mitigates scanner-related confounding.

**Figure 2:**
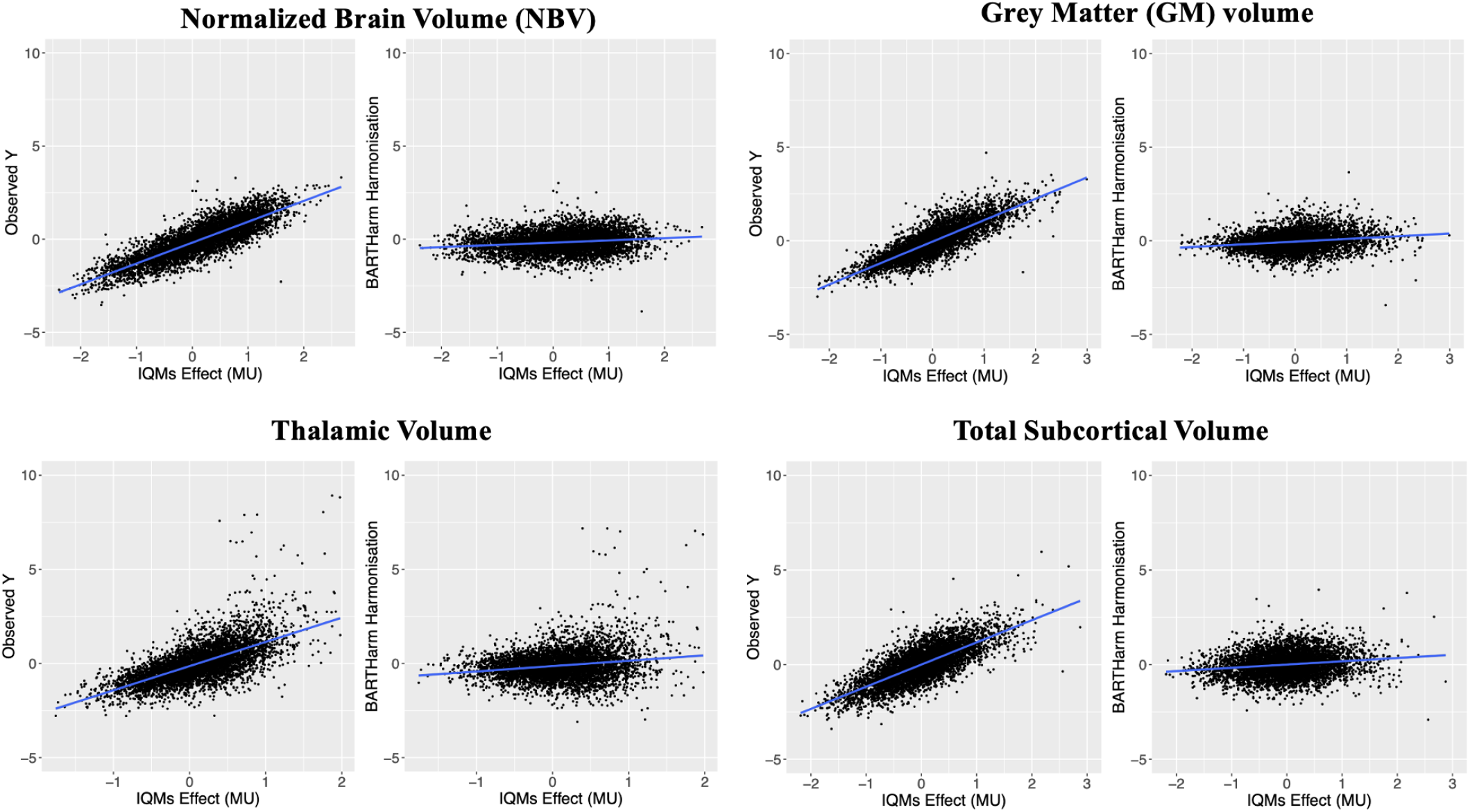
Representative IDPs plotted against the estimated scanner effect (MU *µ*) before and after BARTharm harmonization. Prior to harmonization, a clear trend (i.e., a slope different from zero) indicates residual scanner influence; after BARTharm harmonization, the relationship is effectively eliminated (flattened lines), demonstrating the method’s ability to remove scanner-induced biases. Across all 29 brain IDPs post-harmonization slopes were reduced to the range [0.05, 0.2], representing a statistically significant reduction in scanner-IDP association (*p*-values ≪ 10^−16^).

Finally, to further evaluate whether scanner or site effects were fully removed by BARTharm harmonization, we tested the ability of a support vector machine (SVM) classifier to predict proxy scanner IDs from the harmonized IDPs. Since true scanner labels were not available in NO.MS, we first applied principal component analysis (PCA) to the IQMs to reduce dimensionality and capture the major axes of technical variability. The first 15 principal components explained over 95% of the variance, and the first 10 over 90%. We then applied k-means clustering to the 15 PCA components reduced data to infer scanner-like clusters. To determine the optimal number of clusters, we used the elbow method on the within-cluster sum of squares (WSS). This analysis showed that while WSS dropped steeply for the first few clusters, the reduction decreased considerably from 5 clusters and plateaued around 8 clusters, which we selected as the number of proxy scanner IDs.

Using the 8-clusters assignments as scanner proxies, we trained an SVM classifier with a radial basis function (RBF) kernel (Cortes & Vapnik, 1995) to predict cluster ID from the 29 brain IDPs. A harmonization method that successfully removes scanner-related variation should make these cluster labels harder to predict, resulting in lower SVM classification accuracy. We repeated 10-fold cross-validation 10 times to obtain an average accuracy estimates. For the raw IDPs values, the SVM prediction achieved an average of 44.3% classification accuracy. BARTharm harmonized IDPs instead resulted in a lower average accuracy of 21.2%. To assess whether this performance approached chance levels, we used a permutation-based approach to generate a null distribution of accuracies. This involved randomly permuting the cluster labels and re-training the SVM classifier. The resulting null distribution yielded a mean accuracy of 20.5%, reflecting classification performance expected by chance alone. This indicates that the post-harmonization IDPs are nearly indistinguishable with respect to our “proxy” scanner ID, confirming that BARTharm successfully removed scanner-related signal from the data. We repeated the same analysis with 5-clusters to find raw IDPs accuracy of 52.8%, BARTharm harmonized IDPs accuravy of 25.2%, and null distribution of 23.1%, confirming successful elimination of scanner-related effects.

We then investigated whether BARTharm preserves the biological variability in the data. We assessed the proportion of variation in the 29 brain IDPs that could be explained by three key biological variables, namely Age, Expanded Disability Status Scale (EDSS), and Volume of T2 lesions (VOLT2), before and after BARTharm harmonization. For each IDP and variable, we fit a linear model and computed the coefficient of determination (*R*^2^) as a measure of effect size. To evaluate the change in explanatory power after harmonization, we used Bland–Altman plots, which display the difference in *R*^2^ (post – pre) against the mean *R*^2^ across both conditions. As shown in Figure 3, harmonization consistently increased the variance explained by each biological variable across a majority of IDPs. For age (left panel), several IDPs showed substantial increases in *R*^2^, with a clear upward shift relative to the no-change line, and a strong positive mean difference. A similar but less pronounced pattern was observed for EDSS (middle panel), suggesting that harmonization preserved or modestly enhanced clinical signal. For VOLT2 (right panel), the effect was more variable, but again skewed toward improved explanatory power after harmonization. Together, these results suggest that BARTharm not only removes scanner-related noise, but also enhances the ability of IDPs to reflect biologically meaningful variation.

**Figure 3:**
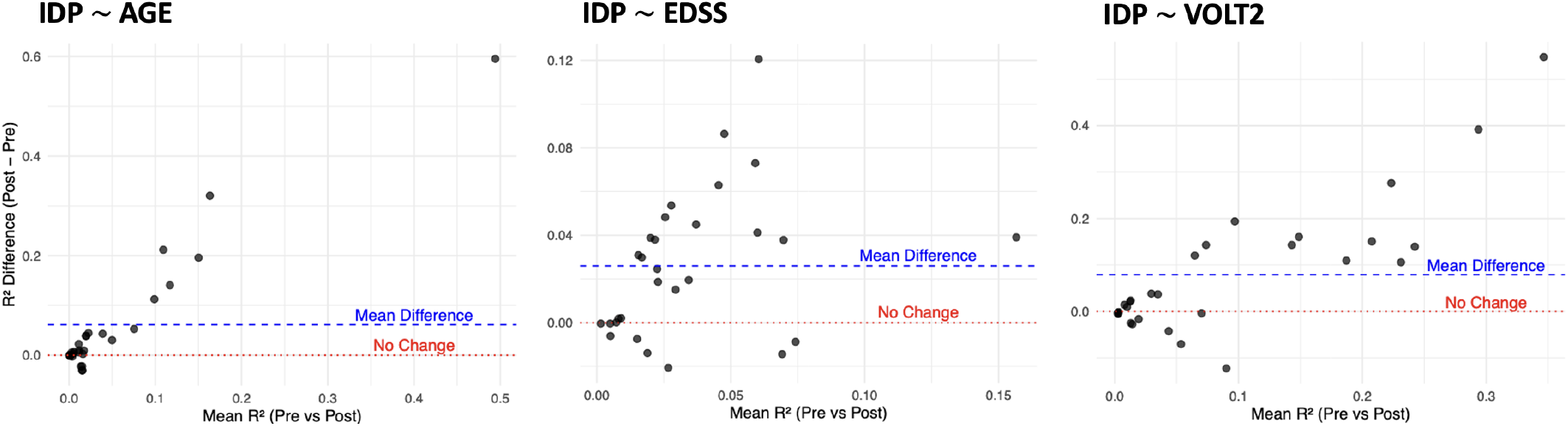
Bland–Altman plots showing the change in *R*^2^ (post – pre) for each IDP (back dots) when regressed on Age (left), EDSS (middle), and VOLT2 (right), plotted against the mean *R*^2^ across pre- and post-harmonization. The red dotted line indicates no change; the blue dashed line indicates the average *R*^2^ difference across IDPs. Most points lie above the red line, suggesting that BARTharm increased the biological signal captured by the IDPs.

## 5 Discussion

Harmonization is essential in multi-site MRI studies to ensure that biological variability can be accurately interpreted without being confounded by scanner-specific effects. Differences in scanner hardware, acquisition protocols, and image quality can introduce systematic differences in imaging-derived phenotypes (IDPs) which becomes particularly problematic when combining data from different trials or large consortia. Without harmonization, downstream analyses can produce biased or misleading results, limiting the translational value of neuroimaging research on large datalakes. As the field increasingly relies on large, heterogeneous datasets, the need for robust, flexible, and generalizable harmonization techniques has become more critical than ever (Laird, 2021; Van Horn & Toga, 2009).

ComBat has emerged as the most widely used statistical harmonization method (Fortin et al., 2017). It assumes that scanner affects are both additive and multiplicative, and estimates these effects for each site using Scanner IDs as categorical labels. ComBat then removes these scanner-induced effects and reintroduces the biological component, preserving variability attributed to known covariates. However, ComBat is inherently limited by its linear modeling assumptions and its reliance on explicit Scanner IDs, which may not be available in anonymized datasets or sufficient to capture subtle inter-scanner differences.

In this paper, we introduced BARTharm, a novel MRI harmonization method designed to address the key limitations of ComBat. Most notably, BARTharm moves away from the reliance on Scanner IDs, acknowledging that these categorical labels are often unavailable, anonymized, or insufficient to capture meaningful inter-scanner variability, and instead uses continuous Image Quality Metrics (IQMs) to model scanner effects (Esteban et al., 2017). These metrics, which are derived directly from the images, provide a more nuanced and quantitative view of scanner-induced variation. In doing so, BARTharm also acknowledges an important practical reality: that even the same scanner, when operating under different protocols or at different times, may require harmonization. At the core of BARTharm is the use of Bayesian Additive Regression Trees (BART) (H. A. Chipman et al., 2010), a non-parametric modeling approach that can flexibly capture complex, non-linear relationships and interactions in the data—both in scanner effects and biological signals. This allows BARTharm to go beyond the linear assumptions of ComBat and adapt to the complex, high-dimensional nature of real-world neuroimaging data. Moreover, by modeling biological and scanner components separately, and by producing full posterior distributions over harmonized outcomes, BARTharm supports rigorous uncertainty quantification, which is essential for reliable downstream inference.

Our simulation studies highlighted the clear advantages of BARTharm across both controlled and realistic settings. In the first simulation framework, based on fully synthetic data, we systematically evaluated performance under different combinations of linear and non-linear scanner and biological effects. Across all scenarios, BARTharm consistently outperformed ComBat, achieving lower RMSE, stronger correlation with the true biological signal, and weaker correlation with scanner-induced noise. Notably, even in the fully linear setting where ComBat is theoretically well-suited, BARTharm still demonstrated improved performance, largely thanks to its use of continuous IQMs rather than categorical Scanner IDs. ComBat relies on scanner IDs to define discrete groups for estimating additive and multiplicative effects, assuming all variability between sites can be captured by a single label. In contrast, IQMs provide a high-dimensional, continuous characterization of scanner-related image quality measurements, capturing subtle differences even within scanners or across protocols used on the same device. As a result, BARTharm is able to exploit richer information about scanner-induced noise, leading to more precise estimation and removal of non-biological effects. Theis shows that, beyond modeling flexibility, the quality of the information used to characterize scanner effects plays a critical role in successful harmonization. When non-linearity was introduced in the biological component, ComBat’s limitations became more evident. Because it assumes a linear relationship between the biological covariates and the imaging outcome, ComBat was unable to model complex, non-linear biological patterns that are commonly observed in neuroimaging data, such as age-related trajectories. This mismatch between model assumptions and the true data-generating process led to inaccurate estimation of the biological signal, and noticeable reduction in the correlation with the true, clean outcome. We also observed that ComBat was more prone to inflated False Positive Rates, indicating a risk of detecting spurious group effects in harmonized outcomes.

In the second simulation framework, using semi-synthetic data constructed from real biological and IQM covariates from the NO.MS dataset, BARTharm continued to perform robustly under more realistic challenges. Here, both the biological and the scanner effects were simulated linearly. In the model misspecification scenario, where only a subset of the true biological covariates was used for harmonization, both methods showed deterioration in performance compared to the similar fully-specified model (Row 1 in Table 1). However, BARTharm was markedly more robust than ComBat: it maintained higher biological correlation and lower residual scanner effects, as well as lower RMSE. In general, we argue that when performing harmonization, it is best to include all available biological covariates of interest in the model. Omitting relevant variables can confound scanner and biological effects, making it more difficult to correctly attribute variability to its true source. This not only reduces the effectiveness of harmonization but can also bias downstream analyses. By incorporating a complete and informative set of covariates, harmonization methods can more accurately separate non-biological scanner effects from meaningful biological variation, ensuring the integrity of subsequent statistical and clinical interpretations.

In the correlation scenario, where scanner assignment was partially confounded with biological effects, BARTharm maintained strong performance, successfully disentangling scanner-related variability from true biological signal. Even under increasing levels of confounding, BARTharm preserved high biological correlation, low scanner residual effects, and controlled false positive rates, highlighting its robustness to challenges frequently encountered in multi-site neuroimaging studies. In contrast, ComBat’s performance deteriorated notably: although it achieved moderately high biological correlations at stronger levels of confounding, it failed to eliminate scanner-driven variability, as reflected by high IQMs correlation, and critically showed substantial inflation of the False Positive Rate (FPR). This result is particularly important because it highlights an important practical limitation of ComBat. Scanner-population confounding is often present in real-world, retrospective, multi-site datasets, where different scanners often scan distinct populations with varying demographics or clinical characteristics. In such scenarios, ComBat can erroneously attribute scanner-related variation to biological differences, as shown in our simulations where ComBat’s FPR substantially exceeded the expected confidence bounds even under moderate confounding. These findings underscore the critical need for more flexible, data-driven harmonization approaches like BARTharm, which can model non-linear, high-dimensional relationships between scanner effects and biological variation without imposing restrictive assumptions. By adapting to the complexity inherent in real-world data, BARTharm demonstrates robustness, flexibility, and reliability in harmonizing MRI data under diverse and practically relevant conditions.

The real-data analysis on the NO.MS dataset highlighted BARTharm’s practical utility in settings where traditional harmonization methods, such as ComBat, cannot be applied due to the unavailability of scanner IDs. In this real-world dataset, BARTharm demonstrated that IQMs can be used to identify and correct for scanner-induced variability, and successfully eliminate scanner-related signal even in the absence of explicit scanner identifiers. Importantly, BARTharm did not just remove noise. The resulting set of harmonized imaging-derived phenotypes (IDPs) exhibited stronger and more biologically plausible associations with clinical outcomes, providing evidence that harmonization improves the clarity and strength of true biological signal. This is a critical requirement for multi-site studies, where the ability to correct non-biological noise without suppressing genuine effects is essential. Moreover, our findings reveal that anonymization does not eliminate the need for harmonization, but rather shifts the challenge toward more data-driven, flexible solutions. These results underscore the limitations of relying solely on Scanner IDs and demonstrate the value of moving toward harmonization frameworks like BARTharm that use rich, intrinsic image information to correct scanner biases while preserving meaningful biological patterns.

BARTharm is not without limitations. While existing harmonization methods such as ComBat operate within a location-scale framework, adjusting both the mean and variance of imaging features across scanners, BARTharm, in its current form, models only scanner-related shifts in the mean (“location” effects). If scanner-related differences in variance (scale effects) are driven by the IQMs, then BARTharm’s flexible, data-driven, non-parametric BART prior can implicitly capture these through non-linear interactions. However, if the source of scale heterogeneity lies outside the IQMs, then the current formulation may fall short. Extending BARTharm to a full location-scale model is a natural next step. Existing heteroscedastic BART implementations, which separately model conditional means and variances (Pratola et al., 2020), provide a clear and promising pathway for this future development. Another limitation relates to computational scalability. Because BARTharm relies on Gibbs sampling and fits each imaging-derived phenotype (IDP) individually, it introduces higher computational overhead compared to simpler models like ComBat. Although parallelization can mitigate some of these costs, further optimization will be essential to enable voxel-level harmonization. Finally, a promising extension would be to adapt BARTharm for longitudinal data settings, drawing inspiration from recent advancements in longitudinal Bayesian Causal Forest models (McJames et al., 2024; Wang et al., 2024; Yeager et al., 2019). A longitudinal version of BARTharm would allow for harmonization across multiple timepoints, helping to disentangle scanner effects from biological changes over time. This would further enrich our ability to track disease progression and treatment response in longitudinal neuroimaging studies, addressing a major current challenge in the field.

## Data and Code Availability

The R code to use BARTharm is publicly available at https://github.com/NeuroSML/BARTharm.

For NO.MS data, the reader is able to request the raw data (anonymized) and related documents (e.g., protocol, reporting and analysis plan, clinical study report) of all the studies that underlie the modelling results reported in this article by connecting to https://www.clinicalstudydatarequest.com and signing a Data Sharing Agreement with Novartis. These will be made available to qualified external researchers, with requests reviewed and approved by an independent review panel on the basis of scientific merit.

## Author Contributions

EP: Conceptualization, Formal analysis, Investigation, Methodology, Software, Validation, Visualization, Writing - original draft, Writing - review & editing. DAH: Resources, Writing - review & editing. LG: Resources, Writing - review & editing. RTS: Writing - review & editing. CCH: Project administration, Supervision. TEN: Project administration, Supervision, Writing - review & editing. HG: Conceptualization, Data curation, Project administration, Supervision, Writing - review & editing.

## Supplementary Material

### 1.1 Mathematical details of Simulation studies

### 1.2 Simulation framework 1

We simulate data for 1,000 subjects under various combinations of linear and non-linear scanner and biological effects. The goal is to assess harmonization performance in increasingly complex scenarios while maintaining a known ground truth. For each subject, we generate covariates, scanner properties, and outcome components as follows:

#### Scanner properties

We define 2 or 10 scanners, each assigned a resolution category and a noise level. From these properties, we derive the signal-to-noise ratio (SNR) and contrast-to-noise ratio (CNR). These quantities are later used to simulate scanner effects and influence measurement noise.

- Scanner Resolution: Each scanner is assigned one of three possible resolutions: *low, medium*, or *high*. The assignment is based on the scanner ID as follows:

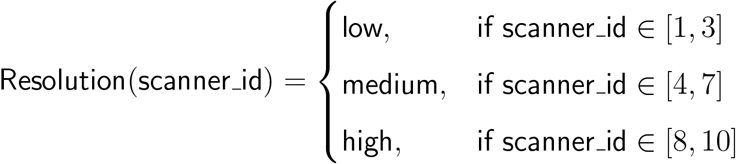

When we instead simulated 2 scanners, we just randomly chose one as high resolution and the other as medium.
- Noise Level: The noise level is inversely related to the scanner ID (higher ID = lower noise):

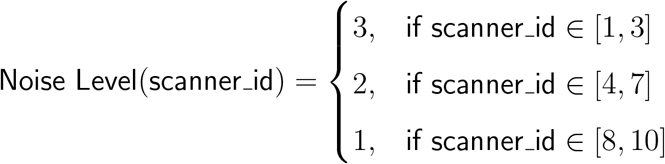

When we instead simulated 2 scanners, we just randomly chose one and assigned a Noise level of 3, and the other of 1.
- Signal-to-Noise Ratio (SNR): SNR measures the signal relative to noise and is based on the scanner resolution, adjusted by noise level:

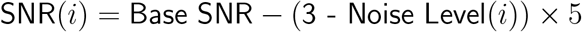

where the Base SNR is generated from Uniform distribution:

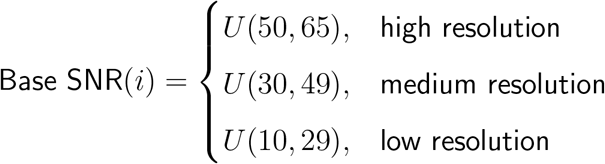
- Contrast-to-Noise Ratio (CNR): CNR measures contrast relative to noise and is similarly affected by resolution and noise level:

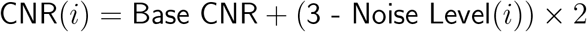

CNR(*i*) = Base CNR + (3 - Noise Level(*i*)) × 2 where the Base CNR is generated from Uniform distribution:

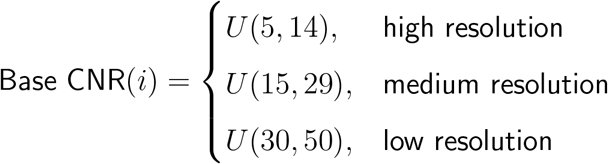

#### Subject-level covariates

We then simulate subject-level data including variables such as age, sex, and scanner ID. For each individual, we simulate:

- **Age:** ∼ 𝒰(20, 60)
- **Sex:** ∼ Bernoulli(0.5)
- **Binary group:** ∼ Bernoulli(0.5)
- **Scanner ID:** Assigned randomly.
- **Biological features:** Two continuous variables, Bio_1_ ∼ 𝒰(0, 15) and Bio_2_ ∼ 𝒰(0, 10)

#### Biological component *Y*_bio_

Depending on the simulation setting, biological effects can be linear (e.g., age, sex, and bio covariates enter additively) or non-linear (e.g., quadratic age, sinusoidal age effects, interactions, and log-transformed bio variables). The binary group has no effect. Gaussian noise is added to simulate within-subject variability.

- **Sex Effect:** Males (sex = 0) have a -25 effect on the outcome, while females (sex = 1) have a +25 effect.
- **Age Effect (Linear Setting):** A negative effect modeled linearly with coefficient -1.5:

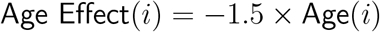
- **Age Effect (Non-linear Setting):** Includes both quadratic and sinusoidal components:

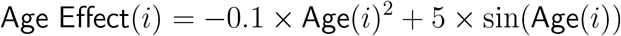
- **Biological Feature Effects (Linear Setting):** Feature 1 has a negative coefficient of -4, while Feature 2 has a positive coefficient of +6:

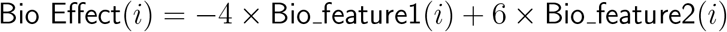
- **Biological Feature Effects (Non-linear Setting):** Includes squared and logarithmic terms:

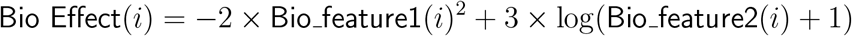
- **Interaction Effects:**

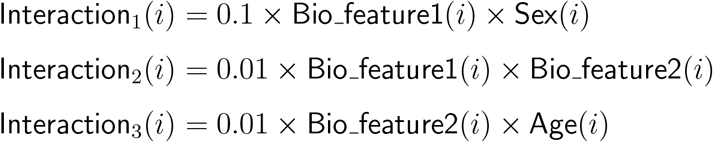
- **Random Effect:** A subject-level noise term is added:

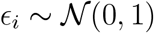

The baseline biological outcome *Y*_bio_ (before scanner effects) is given by:

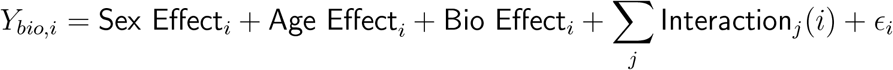

#### Scanner component *Y*_scanner_

Scanner effects are modeled using a combination of resolution, noise level, SNR, and CNR. These effects may be linear (e.g., additive terms and linear interactions) or non-linear (e.g., squared terms, exponential transformations, or interactions between scanner properties). Scanner effects are added to the biological component to form the final outcome.

- **Effect of Scanner Resolution (Linear Setting):** Higher scanner resolution reduces the outcome, modeled linearly:

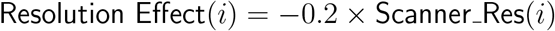
- **Effect of Noise Level (Linear Setting):** Increased noise level raises the outcome:

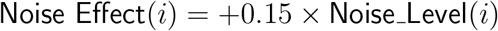
- **Resolution-Noise Interaction (Linear Setting):** Interaction between scanner resolution and noise level:

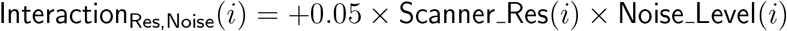
- **SNR and CNR Effects (Linear Setting):** Signal-to-Noise and Contrast-to-Noise Ratios impact the outcome as follows:

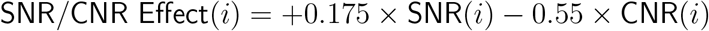
- **SNR-CNR Interaction (Linear Setting):**

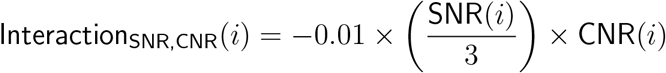
- **Resolution-Noise Effects (Non-linear Setting):** In more complex scenarios, scanner effects are modeled using non-linear transformations of resolution and noise:

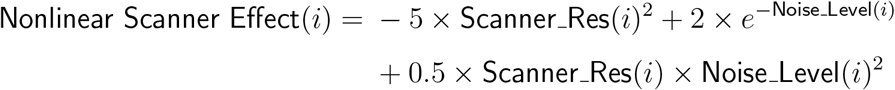
- **SNR and CNR Effects (Non-linear Setting):** In the non-linear case, SNR and CNR contribute as:

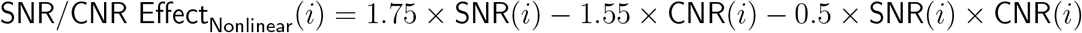

The total scanner-related effect *Y*_*scanner*_ added to the outcome is given by:

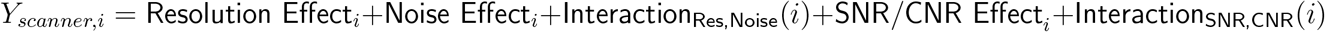

##### 1.2.1 2 Scanners Results

**Table 3:**
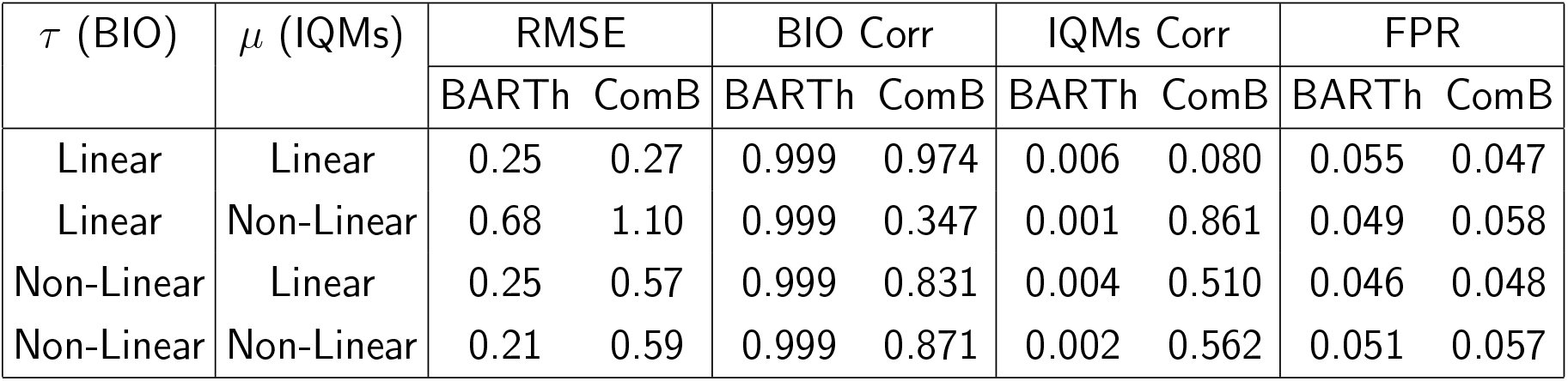
Comparison of BARTharm (BARTh) and ComBat (ComB) under different combinations of linear and non-linear biological (*τ*) and scanner (*µ*) effects, with 2 simulated scanners. RMSE indicates accuracy in recovering the true, harmonized, outcome, BIO Corr measures alignment with the true biological signal, IQMs Corr captures residual scanner effects, FPR denotes the false positive rate in detecting a group effect. Lower RMSE, higher BIO Corr, and lower IQMs Corr collectively signify more effective harmonization.

| $\tau$ (BIO) | $\mu$ (IQMs) | RMSE | | BIO Corr | | IQMs Corr | | FPR | |
| --- | --- | --- | --- | --- | --- | --- | --- | --- | --- |
|  |  | BART<sub>h</sub> | ComB | BART<sub>h</sub> | ComB | BART<sub>h</sub> | ComB | BART<sub>h</sub> | ComB |
| Linear | Linear | 0.25 | 0.27 | 0.999 | 0.974 | 0.006 | 0.080 | 0.055 | 0.047 |
| Linear | Non-Linear | 0.68 | 1.10 | 0.999 | 0.347 | 0.001 | 0.861 | 0.049 | 0.058 |
| Non-Linear | Linear | 0.25 | 0.57 | 0.999 | 0.831 | 0.004 | 0.510 | 0.046 | 0.048 |
| Non-Linear | Non-Linear | 0.21 | 0.59 | 0.999 | 0.871 | 0.002 | 0.562 | 0.051 | 0.057 |

### 1.3 Simulation framework 2

In the second simulation framework, we use real data from NO.MS dataset, including real Image Quality Metrics (IQMs) *X*_*nonbio*_ and real biological covariates *X*_*bio*_. We then simulated Scanner IDs, regression coefficients for both *X*_*nonbio*_ and *X*_*bio*_, and the final outcome to be harmonized *Y*_*sim*_.

#### Scanner ID Simulation

To generate scanner IDs, we apply *k*-means clustering to the normalized IQM covariates and assign each subject to one of five clusters. These cluster labels serve as proxy Scanner IDs and introduce structured scanner-related variability.

#### Biological Effect *Y*_bio_

We sample a set of regression coefficients *β*_bio_ ∼ 𝒩 (2, 4), which are constant across Scanner IDs, and apply them to the biological covariates matrix *X*_bio_ to generate the true biological signal:

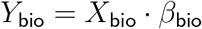

#### Scanner Effect *Y*_scanner_

To simulate scanner-specific effects, we first generate a baseline coefficient vector *β*_nonbio_ ∼ 𝒩 (0, 0.5) for the non-biological covariates. Then, for each scanner cluster, we perturb *β*_nonbio_ by adding scanner-specific Gaussian noise with scanner-dependent variances. Specifically, for each scanner *s*, the scanner-specific coefficients 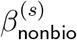 are sampled as:

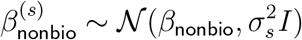

where 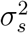 is the scanner-specific variance chosen from a predefined set {0.2, 0, 0.5, 1, 2} depending on the scanner ID. Each subject *i* is assigned the corresponding scanner-specific coefficient vector based on their simulated scanner ID. Their scanner-related outcome contribution is then computed as:

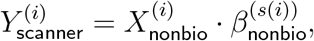

where *s*(*i*) denotes the scanner assignment for subject *i*. This modeling approach induces scanner-specific heterogeneity in the effect of non-biological covariates, realistically simulating variability commonly observed across sites.

#### Introducing correlation between *Y*_scanner_ and *Y*_bio_

In settings where we aim to simulate correlation (i.e., confounding) between biological and scanner effects, we modify the construction of *β*_nonbio_ to partially depend on *β*_bio_. Specifically, we start by copying the coefficients from *β*_bio_ into *β*_nonbio_, cyclically repeat elements of *β*_bio_ to fill *β*_nonbio_, thus inducing similarity between biological and scanner-related coefficients at the feature level. Then, for each scanner *s*, we generate scanner-specific coefficients by scaling *β*_nonbio_ with a predefined scanner-dependent multiplier:

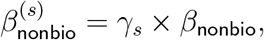

where *γ*_*s*_ ∈ {−1, −0.5, 0.5, 1, 2} is a fixed scalar associated with each scanner ID. This scaling step introduces systematic variation between scanners while preserving an underlying correlation structure with the biological signal. Each subject *i* is assigned the corresponding scanner-specific coefficient vector based on their simulated scanner ID. Their scanner-related outcome contribution is then computed as:

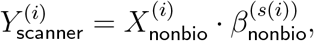

where *s*(*i*) denotes the scanner assignment for subject *i*.

The resulting *Y*_scanner_ and *Y*_bio_ are then combined in a controlled fashion to achieve a target correlation (e.g., 0.30, 0.50, 0.75) between the two components:

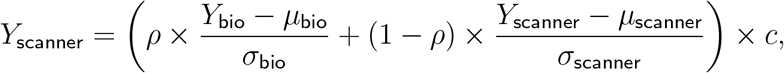

where *µ* and *σ* are the mean and standard deviation of each component, and *c* and *ρ* are scaling constant which are tuned at the beginning to achieve the target correlation ±5 in each realization of the data.

#### Total Outcome

The simulated outcome is:

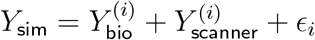

where *ϵ*_*i*_ ∼ 𝒩 (0, *σ*^2^) is Gaussian noise scaled to 5% of the total outcome standard deviation. *Y*_clean_ which is the target, harmonised, outcome, is given by 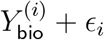.

#### Group Confounding

To simulate realistic confounding, we introduce a binary group variable based on the IQM most strongly associated with scanner effects. We identify the IQM with the largest average absolute effect (from *β*_scanner_) and bin subjects into “low” and “high” IQM groups based on a threshold (0.5 as the data is quantile normalized). Within each bin, subjects are randomly assigned to group 0 or 1 with bin-specific probabilities (e.g., 45% vs. 55%) which are chosen ensuring that we are not inadvertently and indirectly introducing any effect on *Y*_bio_, especially when *Y*_bio_ and *Y*_scanner_ are correlated.

This creates a weak but controlled association between scanner-induced variability and group assignment. It follows that the group assignment will have a non-zero effect on the simulated outcome *Y*_*sim*_ but no effect on *Y*_bio_ as the *β*_*bio*_ for the group variable is zero.

### 1.4 NO.MS IDPs harmonization

The full list of harmonized IDPs is the following:

- Normalized Brain Volume (NBV)
- Total Subcortical volume (includes (Thalamus, Caudate, Pallidum, Putamen, Brainstem, Hippocampus, Amygdala, Accumbens)
- Ventricular Volume
- Brainstem Volume
- Grey Matter Volume
- MNI Cerebellum
- MNI Frontal
- MNI Caudate
- MNI Putamen
- Insula
- Temporal
- Parietal
- Occipital
- Thalamus, Left Thalamus and Right Thalamus
- Left Caudate and Right Caudate
- Left Putamen and Right Putamen
- Left Pallidum and Right Pallidum
- Left Amygdala and Right Amygdala
- Left Accumbens and Right Accumbens
- Left Hippocampus and Right Hippocampus

## Footnotes

1 We ran similar experiments where we randomly selected 50% of the biological covariates and kept the selection fixed across all 1000 realizations and observed very minimal difference in performance from both ComBat and BARTharm.

## References

Acquitter, C., Piram, L., Sabatini, U., Gilhodes, J., Moyal Cohen-Jonathan, E., Ken, S., & Lemasson, B. (2022). Radiomics-based detection of radionecrosis using harmonized multiparametric mri. Cancers, 14 (2), 286.

An, L., Zhang, C., Wulan, N., Zhang, S., Chen, P., Ji, F., Ng, K. K., Chen, C., Zhou, J. H., Yeo, B. T., et al. (2024). Deepresbat: Deep residual batch harmonization accounting for covariate distribution differences. Medical Image Analysis, 103354.

Barth, C., Kelly, S., Nerland, S., Jahanshad, N., Alloza, C., Ambrogi, S., Andreassen, O. A., Andreou, D., Arango, C., Baeza, I., et al. (2023). In vivo white matter microstructure in adolescents with early-onset psychosis: A multi-site mega-analysis. Molecular Psychiatry, 28 (3), 1159–1169.

Bayer, J. M., Thompson, P. M., Ching, C. R., Liu, M., Chen, A., Panzenhagen, A. C., Jahanshad, N., Marquand, A., Schmaal, L., & Sämann, P. G. (2022). Site effects how-to and when: An overview of retrospective techniques to accommodate site effects in multi-site neuroimaging analyses. Frontiers in neurology, 13, 923988.

Beer, J. C., Tustison, N. J., Cook, P. A., Davatzikos, C., Sheline, Y. I., Shinohara, R. T., Linn, K. A., Initiative, A. D. N., et al. (2020). Longitudinal combat: A method for harmonizing longitudinal multi-scanner imaging data. Neuroimage, 220, 117129.

Campello, V. M., Martin-Isla, C., Izquierdo, C., Guala, A., Palomares, J. F. R., Viladés, D., Descalzo, M. L., Karakas, M., Çavuş, E., Raisi-Estabragh, Z., et al. (2022). Minimising multi-centre radiomics variability through image normalisation: A pilot study. Scientific reports, 12 (1), 12532.

Chen, A. A., Beer, J. C., Tustison, N. J., Cook, P. A., Shinohara, R. T., Shou, H., & Initiative, A. D. N. (2022). Mitigating site effects in covariance for machine learning in neuroimaging data. Human brain mapping, 43 (4), 1179–1195.

Chipman, H., George, E., & McCulloch, R. (2006). Bayesian ensemble learning. Advances in neural information processing systems, 19.

Chipman, H. A., George, E. I., & McCulloch, R. E. (2010). Bart: Bayesian additive regression trees.

Clark, K. A., Woods, R. P., Rottenberg, D. A., Toga, A. W., & Mazziotta, J. C. (2006). Impact of acquisition protocols and processing streams on tissue segmentation of t1 weighted mr images. NeuroImage, 29 (1), 185–202.

Cortes, C., & Vapnik, V. (1995). Support-vector networks. Machine learning, 20, 273–297.

Dahlke, F., Arnold, D. L., Aarden, P., Ganjgahi, H., Häring, D. A., Čuklina, J., Nichols, T. E., Gardiner, S., Bermel, R., & Wiendl, H. (2021). Characterisation of ms phenotypes across the age span using a novel data set integrating 34 clinical trials (no. ms cohort): Age is a key contributor to presentation. Multiple Sclerosis Journal, 27 (13), 2062–2076.

Dai, P., Xiong, T., Zhou, X., Ou, Y., Li, Y., Kui, X., Chen, Z., Zou, B., Li, W., Huang, Z., et al. (2022). The alterations of brain functional connectivity networks in major depressive disorder detected by machine learning through multisite rs-fmri data. Behavioural Brain Research, 435, 114058.

Esteban, O., Birman, D., Schaer, M., Koyejo, O. O., Poldrack, R. A., & Gorgolewski, K. J. (2017). Mriqc: Advancing the automatic prediction of image quality in mri from unseen sites. PloS one, 12 (9), e0184661.

Esteban, O., Blair, R. W., Nielson, D. M., Varada, J. C., Marrett, S., Thomas, A. G., Poldrack, R. A., & Gorgolewski, K. J. (2019). Crowdsourced mri quality metrics and expert quality annotations for training of humans and machines. Scientific data, 6 (1), 30.

Fjell, A. M., Walhovd, K. B., Westlye, L. T., Østby, Y., Tamnes, C. K., Jernigan, T. L., Gamst, A., & Dale, A. M. (2010). When does brain aging accelerate? dangers of quadratic fits in cross-sectional studies. Neuroimage, 50 (4), 1376–1383.

Fortin, J.-P., Parker, D., Tunç, B., Watanabe, T., Elliott, M. A., Ruparel, K., Roalf, D. R., Satterthwaite, T. D., Gur, R. C., Gur, R. E., et al. (2017). Harmonization of multi-site diffusion tensor imaging data. Neuroimage, 161, 149–170.

Garcia-Dias, R., Scarpazza, C., Baecker, L., Vieira, S., Pinaya, W. H., Corvin, A., Redolfi, A., Nelson, B., Crespo-Facorro, B., McDonald, C., et al. (2020). Neuroharmony: A new tool for harmonizing volumetric mri data from unseen scanners. Neuroimage, 220, 117127.

Goto, M., Abe, O., Miyati, T., Kabasawa, H., Takao, H., Hayashi, N., Kurosu, T., Iwatsubo, T., Yamashita, F., Matsuda, H., et al. (2012). Influence of signal intensity non-uniformity on brain volumetry using an atlas-based method. Korean Journal of Radiology, 13 (4), 391.

Hahn, P. R., Carvalho, C. M., Puelz, D., & He, J. (2018). Regularization and confounding in linear regression for treatment effect estimation.

Hahn, P. R., Murray, J. S., & Carvalho, C. M. (2020). Bayesian regression tree models for causal inference: Regularization, confounding, and heterogeneous effects (with discussion). Bayesian Analysis, 15 (3), 965–1056.

Han, X., Jovicich, J., Salat, D., van der Kouwe, A., Quinn, B., Czanner, S., Busa, E., Pacheco, J., Albert, M., Killiany, R., et al. (2006). Reliability of mri-derived measurements of human cerebral cortical thickness: The effects of field strength, scanner upgrade and manufacturer. Neuroimage, 32 (1), 180–194.

Hill, J., Linero, A., & Murray, J. (2020). Bayesian additive regression trees: A review and look forward. Annual Review of Statistics and Its Application, 7, 251–278.

Hu, F., Chen, A. A., Horng, H., Bashyam, V., Davatzikos, C., Alexander-Bloch, A., Li, M., Shou, H., Satterthwaite, T. D., Yu, M., et al. (2023). Image harmonization: A review of statistical and deep learning methods for removing batch effects and evaluation metrics for effective harmonization. NeuroImage, 274, 120125.

Ingalhalikar, M., Shinde, S., Karmarkar, A., Rajan, A., Rangaprakash, D., & Deshpande, G. (2021). Functional connectivity-based prediction of autism on site harmonized abide dataset. IEEE transactions on biomedical engineering, 68 (12), 3628–3637.

Johnson, W. E., Li, C., & Rabinovic, A. (2007). Adjusting batch effects in microarray expression data using empirical bayes methods. Biostatistics, 8 (1), 118–127.

Jovicich, J., Czanner, S., Greve, D., Haley, E., van Der Kouwe, A., Gollub, R., Kennedy, D., Schmitt, F., Brown, G., MacFall, J., et al. (2006). Reliability in multi-site structural mri studies: Effects of gradient non-linearity correction on phantom and human data. Neuroimage, 30 (2), 436–443.

Laird, A. (2021). Large, open datasets for human connectomics research: Considerations for reproducible and responsible data use. neuroimage, 244, article 118579.

Linero, A. R. (2022). Softbart: Soft bayesian additive regression trees. arXiv preprint 2210.16375.

Mallon, A.-M., Häring, D. A., Dahlke, F., Aarden, P., Afyouni, S., Delbarre, D., El Emam, K., Ganjgahi, H., Gardiner, S., Kwok, C. H., et al. (2021). Advancing data science in drug development through an innovative computational framework for data sharing and statistical analysis. BMC Medical Research Methodology, 21, 1–11.

McJames, N., O’Shea, A., & Parnell, A. (2024). Bayesian causal forests for longitudinal data: Assessing the impact of part-time work on growth in high school mathematics achievement. arXiv preprint 2407.11927.

Moyer, D., Ver Steeg, G., Tax, C. M., & Thompson, P. M. (2020). Scanner invariant representations for diffusion mri harmonization. Magnetic resonance in medicine, 84 (4), 2174–2189.

Pagani, E., Storelli, L., Pantano, P., Petsas, N., Tedeschi, G., Gallo, A., De Stefano, N., Battaglini, M., Rocca, M. A., Filippi, M., et al. (2023). Multicenter data harmonization for regional brain atrophy and application in multiple sclerosis. Journal of Neurology, 270 (1), 446–459.

Pomponio, R., Erus, G., Habes, M., Doshi, J., Srinivasan, D., Mamourian, E., Bashyam, V., Nasrallah, I. M., Satterthwaite, T. D., Fan, Y., et al. (2020). Harmonization of large mri datasets for the analysis of brain imaging patterns throughout the lifespan. NeuroImage, 208, 116450.

Pratola, M. T., Chipman, H. A., George, E. I., & McCulloch, R. E. (2020). Heteroscedastic bart via multiplicative regression trees. Journal of Computational and Graphical Statistics, 29 (2), 405– 417.

Radua, J., Vieta, E., Shinohara, R., Kochunov, P., Quidé, Y., Green, M. J., Weickert, C. S., Weickert, T., Bruggemann, J., Kircher, T., et al. (2020). Increased power by harmonizing structural mri site differences with the combat batch adjustment method in enigma. Neuroimage, 218, 116956.

Ran, H., & Bai, Y. (2021). On soft bayesian additive regression trees and asynchronous longitudinal regression analysis. arXiv preprint 2108.11603.

Reynolds, M., Chaudhary, T., Torbati, M. E., Tudorascu, D. L., Batmanghelich, K., Initiative, A. D. N., et al. (2023). Combat harmonization: Empirical bayes versus fully bayes approaches. NeuroImage: Clinical, 39, 103472.

Sohn, K., Lee, H., & Yan, X. (2015). Learning structured output representation using deep conditional generative models. Advances in neural information processing systems, 28.

Takao, H., Hayashi, N., & Ohtomo, K. (2011). Effect of scanner in longitudinal studies of brain volume changes. Journal of Magnetic Resonance Imaging, 34 (2), 438–444.

Tan, Y. V., & Roy, J. (2019). Bayesian additive regression trees and the general bart model. Statistics in medicine, 38 (25), 5048–5069.

Tang, H., Xu, D., Sebe, N., & Yan, Y. (2019). Attention-guided generative adversarial networks for unsupervised image-to-image translation. 2019 International Joint Conference on Neural Networks (IJCNN), 1–8.

Tardif, C. L., Collins, D. L., & Pike, G. B. (2009). Sensitivity of voxel-based morphometry analysis to choice of imaging protocol at 3 t. Neuroimage, 44 (3), 827–838.

Trefler, A., Sadeghi, N., Thomas, A. G., Pierpaoli, C., Baker, C. I., & Thomas, C. (2016). Impact of timeof-day on brain morphometric measures derived from t1-weighted magnetic resonance imaging. Neuroimage, 133, 41–52.

Van Horn, J. D., & Toga, A. W. (2009). Multisite neuroimaging trials. Current opinion in neurology, 22 (4), 370–378.

Wang, M., Martinez, I., & Hahn, P. R. (2024). Longbet: Heterogeneous treatment effect estimation in panel data. arXiv preprint 2406.02530.

Yeager, D. S., Hanselman, P., Walton, G. M., Murray, J. S., Crosnoe, R., Muller, C., Tipton, E., Schneider, B., Hulleman, C. S., Hinojosa, C. P., et al. (2019). A national experiment reveals where a growth mindset improves achievement. Nature, 573 (7774), 364–369.

Zhu, J.-Y., Park, T., Isola, P., & Efros, A. A. (2017). Unpaired image-to-image translation using cycle-consistent adversarial networks. Proceedings of the IEEE international conference on computer vision, 2223–2232.

